# Convergent antigenic drift of the influenza hemagglutinin lateral patch across time and species

**DOI:** 10.64898/2026.04.16.718966

**Authors:** Jordan I. Lemus-Reyes, Monica L. Fernandez-Quintero, Edgardo Ayala, Olivia M. Swanson, Marina R. Good, Wei Ji, Devika Suja, Julianna Han, Andrew B. Ward, Jenna J. Guthmiller

## Abstract

The lateral patch epitope of the H1 hemagglutinin (HA) was a dominant target of antibodies following exposure to the 2009 pandemic H1N1 virus. However, the conservation and potential for antigenic drift in the lateral patch remain unresolved. Here, we used lateral patch-specific monoclonal antibodies (mAbs) to understand the antigenicity of the lateral patch of human, avian, and swine H1N*x* viruses spanning from 1918 to 2022. We identified discrete mutations that evaded lateral patch-targeting mAbs in pre- and post-2009 H1N1 viruses, leading to genetic differences in lateral patch-targeting antibodies in individuals across birth years. We observed that the lateral patch remains well conserved across zoonotic sources, suggesting existing lateral patch antibodies could protect against a future H1N*x* pandemic. Together, these data support that lateral patch antigenic drift has shaped the human B cell repertoire against influenza viruses and that the lateral patch remains an attractive target for pandemic preparedness.

## Introduction

Influenza A viruses (IAVs) are one of the most common infectious respiratory diseases that threaten the human population. Estimates of seasonal influenza-associated mortality range from 290,000 to 645,000 deaths annually, and more than 5 million influenza-associated hospitalizations occur each year (1,2). Beyond seasonal epidemics, IAVs remain a tangible pandemic threat given their propensity for interspecies transmission, infecting a broad range of avian and mammalian hosts, including swine (3,4). This leads to zoonotic spillover events that can generate reassortant viruses in multiple species and subsequent transmission to humans.

These IAV spillovers have led to four pandemics, two of which were caused by the H1N1 subtype in 1918 and 2009 of avian and swine origins, respectively (5–7). Current efforts to curb the endemic spread of seasonal IAV subtypes, H1N1 and H3N2, are largely focused on annual vaccination efforts. However, influenza viruses rapidly acquire mutations, a process known as antigenic drift, to evade neutralizing antibody responses elicited by past natural infections or vaccinations, prompting the annual reformulation of the influenza vaccine (8).

Antibody responses primarily target the globular head domain of the surface glycoprotein hemagglutinin (HA). HA has five major antigenic sites, which are the main targets of antibodies induced by infection and vaccination (9). These antigenic sites are highly variable over time, across clade lineages, and across geographic regions, resulting in an increasingly genetically and antigenically divergent pool of IAVs(10). Despite this antigenic variability, potent and broadly neutralizing antibodies can target conserved epitopes on the HA head domain, specifically the receptor-binding site (RBS) (11) and the lateral patch (12–15). Previous work has demonstrated that humans induce robust antibody responses against the lateral patch following vaccination with the 2009 pandemic H1N1 (pH1N1) virus and can neutralize pre- and post-2009 H1N1 viruses (12–15). However, a focused antibody response against the lateral patch led to antigenic drift during the 2013-2014 influenza season (16), resulting in a seasonal influenza virus outbreak with greater morbidity and mortality than the 2009 pandemic, particularly among middle-aged adults (17,18). Despite its rapid antigenic drift, the lateral patch epitope remained well-conserved between 1977 and 2009 (12–16). The conservation of the lateral patch across time and animal reservoirs remains poorly investigated and ill-defined.

Here, we utilized characterized lateral patch-targeting mAbs isolated from human donors (12,14) to assess the conservation of the lateral patch across historical H1N*x* isolates from different hosts. We assessed lateral patch-specific mAb binding affinity to H1 HA antigens from 1918-2022 to understand the conservation of the lateral patch in seasonal H1N1 (sH1N1; 1918-2008), pH1N1 (post-2009), and zoonotic H1N*x* isolates. We identified two classes of lateral patch-targeting mAbs based on their antibody gene usage and angle of approach to the lateral patch (12). These distinct antibody classes were linked to donor age; class 1 lateral patch-specific mAbs were found in individuals born post-1980, whereas class 2 lateral patch-specific mAbs were found in individuals born before 1977. We identified critical mutations in HA that have emerged and persisted in post-2009 pH1N1 that can readily escape mAb binding. Despite this, we found one lateral patch-specific mAb that maintains binding to current pH1N1 viruses, suggesting the lateral patch is still an attractive target for generating targeted humoral immune responses. We identified mutations in sH1N1 in the late 1950s and 1970s, when sH1N1 went extinct and was reintroduced into the human population in 1977. These mutations ablated class 1 lateral patch-specific mAb binding, which could explain the birth-year-specific patterns observed in the distinct lateral patch-binding mAb classes. Finally, we found that most lateral patch-targeting mAbs bind to zoonotic H1N*x* antigens, suggesting that the lateral patch epitope is well conserved and could be targeted to mediate protection in the event of zoonotic H1N*x* spillovers. Together, these data support that the lateral patch remains an attractive target for eliciting antibody responses to mediate protection against circulating H1N1 viruses and pandemic H1Nx viruses.

## Results

### Evolution of H1 HA Across Time and Species

H1N1 viruses have circulated in humans since the 1918 pandemic, originating from an ancestral avian H1N1 virus (19) (Fig. 1A). The 1918-lineage seasonal H1N1 (sH1N1) circulated from 1918 to 1957, before being replaced in 1957 by H2N2 (20). Eventually, sH1N1 viruses resembling those from the 1950s reemerged in 1977 and persisted in humans until 2009 (21,22). sH1N1 viruses went extinct in 2009 due to the emergence of the 2009 pH1N1 virus, which derived its HA from the classical swine lineage 1A (7). In parallel, swine H1N*x* viruses have remained in circulation since the 1918 pandemic and have caused occasional swine-to-human influenza infections, including the swine-origin H1N1 influenza outbreak at Fort Dix, New Jersey in 1976 (24). In 1998, a new triple reassortant virus emerged, generating the H1N1 swine virus, which ultimately led to the 2009 pH1N1 pandemic (25). Drifted variants of pH1N1 continue to circulate in humans and cause seasonal outbreaks.

**Figure 1.**
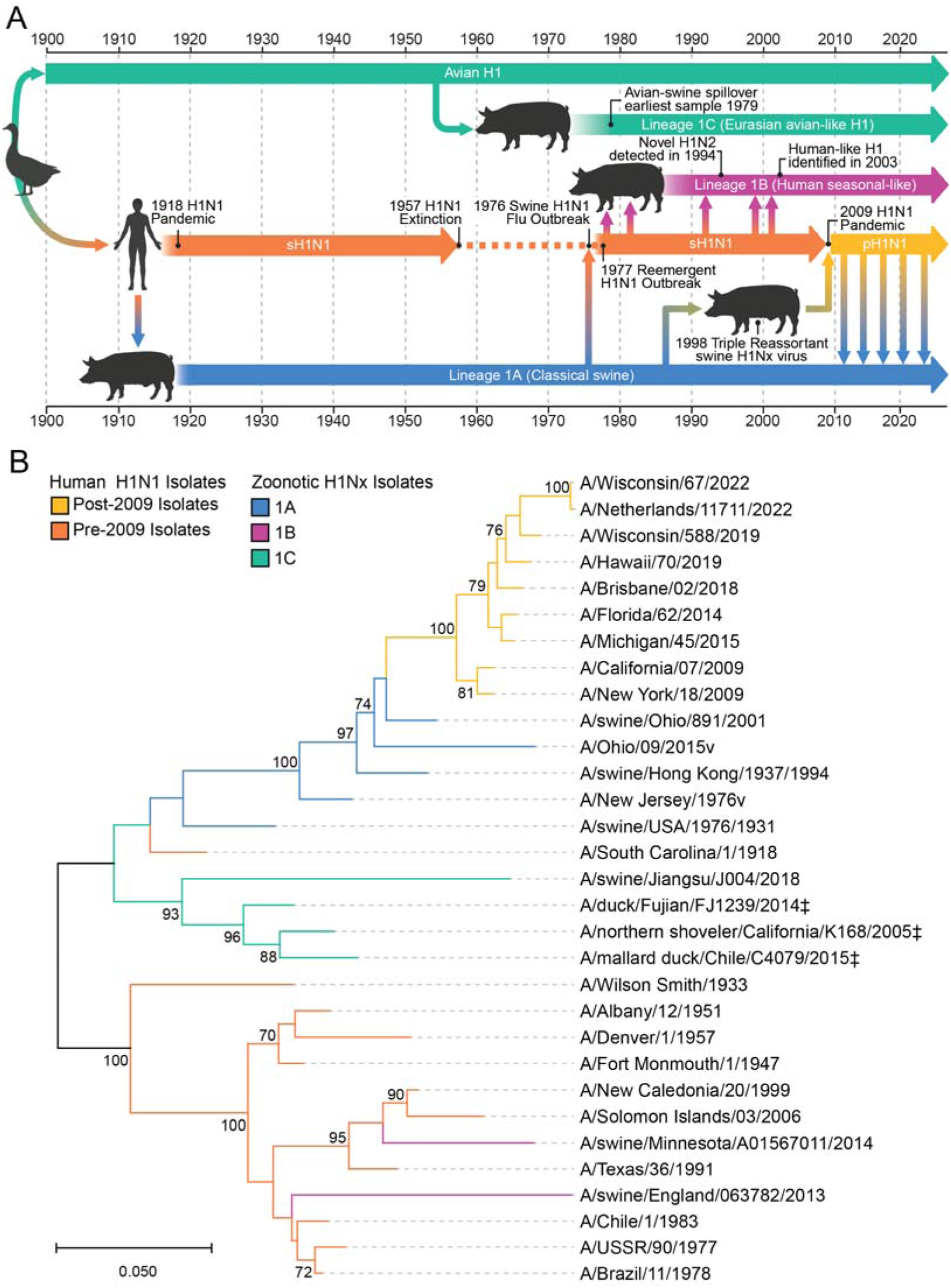
Genetic and phylogenetic relationship of the H1 HA influenza virus in human and zoonotic isolates based on the amino acid sequence. (A) Timeline of human, swine, and avian H1 lineages. (B) Analysis of 31 HA sequences based on the best neighbor-joining method was used to analyze 31 HA sequences from this study (see Table S1). Branch colors are used to distinguish pre- and post-2009 human H1N1 isolates and zoonotic isolates across lineages. The scale bar indicates the branch length corresponding to 0.050 amino acid substitutions per site using the Jones-Taylor-Thornton (JTT) matrix-based method. ‡Avian H1s are marked as 1C lineage, as no clade classification system exists for avian H1N*x* viruses.

Avian and swine H1 viruses co-circulate with human H1N1 viruses (Fig. 1A), with three swine lineages deriving from distinct origins (26). The 1A classical swine lineage is closely related to the 1918 H1N1 pandemic virus (19) and circulates globally, particularly subclade 1A.3.3.2 (27,28). This subclade was the result of reverse zoonosis of the 2009 pH1N1 lineage in pigs, a frequent event that still occurs with contemporary pH1N1 viruses (29–31). Similarly, the 1B swine lineage is derived from multiple, separate events of reverse zoonosis of human sH1N1 viruses between 1977 and 2009. The first 1B clade spillover, initially detected in 1994 in Great Britain, and later spread throughout Europe, likely originated from a circulating sH1N1 in the early 1980s (32,33). Two separate introductions of sH1N1 into swine in the early 2000s were first detected in the US in 2005 (Fig. 1A and 1B) (34–36). These independent spillovers correspond to two distinct 1B clades, each associated with different geographical regions: 1B.1 H1N*x* is most prevalent in the United Kingdom and Europe (26,32,37), whereas 1B.2 H1N*x* predominantly circulates in the Americas and Asia (26,35). Despite these two distinct clades, there is phylogenetic evidence suggesting human seasonal-like H1N1 viruses infected swine populations as early as the 1970s, 1990s, and 2000s in separate geographical regions (38) (Fig. 1A). Lastly, the 1C swine lineage, or the Eurasian avian (EA)-like H1 lineage, is derived from avian H1N*x* viruses and entered the swine population in the 1970s, first appearing in Europe and subsequently spreading throughout the Eurasian continents (34,39) (Fig. 1A). Avian H1s used in this study are classified as 1C lineage given their similarity to swine 1C on sequence-based analysis and because no clade classification system exists for circulating avian H1N*x* viruses (26,40) (Fig. 1B).

To understand the relationship between human and zoonotic H1s, we generated a phylogenetic tree of H1 amino acid sequences derived from sH1N1, pH1N1, the three H1 swine lineages, and the avian H1s related to swine 1C viruses. The joint-joining phylogenetic tree was inferred using all representative H1 HA amino acid sequences (Table S1) to reveal the sequence diversity of known circulating isolates collected over time, from differing host origins, and across various geographic regions. Lineages and clade designations are shown, associated with their corresponding branch colors (Fig. 1A and 1B). 1A and 1B swine H1 isolates cluster near human isolates, indicating the role of reverse zoonosis in their origin. Isolates such as A/New Jersey/1976v (NJ/76v) and A/Ohio/09/2015v (OH/15v) are variant isolates that result from swine-to-human spillovers and are atypical of human seasonal influenza viruses. The HA amino acid segment showed bootstrap support of 70% or more in 1000 replicates, as indicated by the nodes, particularly in most human isolates and a few zoonotic isolates (Fig. 1B). The A/South Carolina/1/1918 (SC/18) virus established the classical swine lineage (1A), which subsequently gave rise to the pH1N1 viruses. These data support that the reverse zoonosis of a 1918-like H1N1 virus in pigs drifted slowly due to the close phylogenetic relatedness of these strains.

SC/18 was also distinct from other post-1918 sH1N1 viruses, which exhibit substantial amino acid diversity from the 1918 isolate. However, the accuracy of older strain sequences (1930s-1950s) remains uncertain, as these viruses were serially passaged in embryonated chicken eggs prior to sequencing, introducing mutations not observed in circulating strains (43,44). sH1N1 viruses continued to undergo antigenic drift until they went extinct in 2009 due to the emergence of the pH1N1 virus.

Despite their genetic differences, zoonotic isolates have limited genetic diversity, specifically in lineages 1A and 1C. This lack of diversity could suggest these viruses undergo slower antigenic drift relative to human H1N1 viruses, potentially as their longevity of these animals’ lifespan limits the immune pressure placed on zoonotic-origin H1 viruses (45,46) (Fig. 1B). However, increase of genetic and antigenic diversity of H1s is caused by intensive farming practices, including indoor housing, transportation within and between farms, naïve and herd-level immunity, and interactions between various species of IAV-susceptible hosts (4,47). While there may have been a greater divergence of amino acid substitutions acquired over time in zoonotic H1s (Fig. 1B), this may be attributed to substantiated gaps in zoonotic H1 surveillance (48). In addition, genomic surveillance and tracking of swine IAVs are limited and challenging due to limited resources among geographic regions (49,50). Together, these data support three major branches of H1 diversity: 1) H1 sequences derived from the initial 1918 spillover into humans and the classical swine lineage (1A), 2) viruses deriving from humans to swine (1B) from 1977-2009, and 3) the avian and EA-like swine lineages of H1 (1C).

### Rapid Antigenic Drift in the Lateral Patch of Post-2009 H1N1 Viruses

Since their introduction in 2009, pH1N1 viruses have undergone significant antigenic drift (51,52). Between 2010 to 2020, 29 amino acid substitutions emerged within in the HA head domain (Table S2); 11 mutations occurred within the variable antigenic sites (Ca1, Ca2, Cb, Sa, and Sb), with one mutation introducing an N-glycosylation motif at position 165 (Fig. 2A). The lateral patch epitope lies in-between the Ca1, Cb, Sa antigenic sites, where a total of 5 mutations were acquired in and around the lateral patch in drifted pH1N1 viruses (Fig. 2A-B).

**Figure 2.**
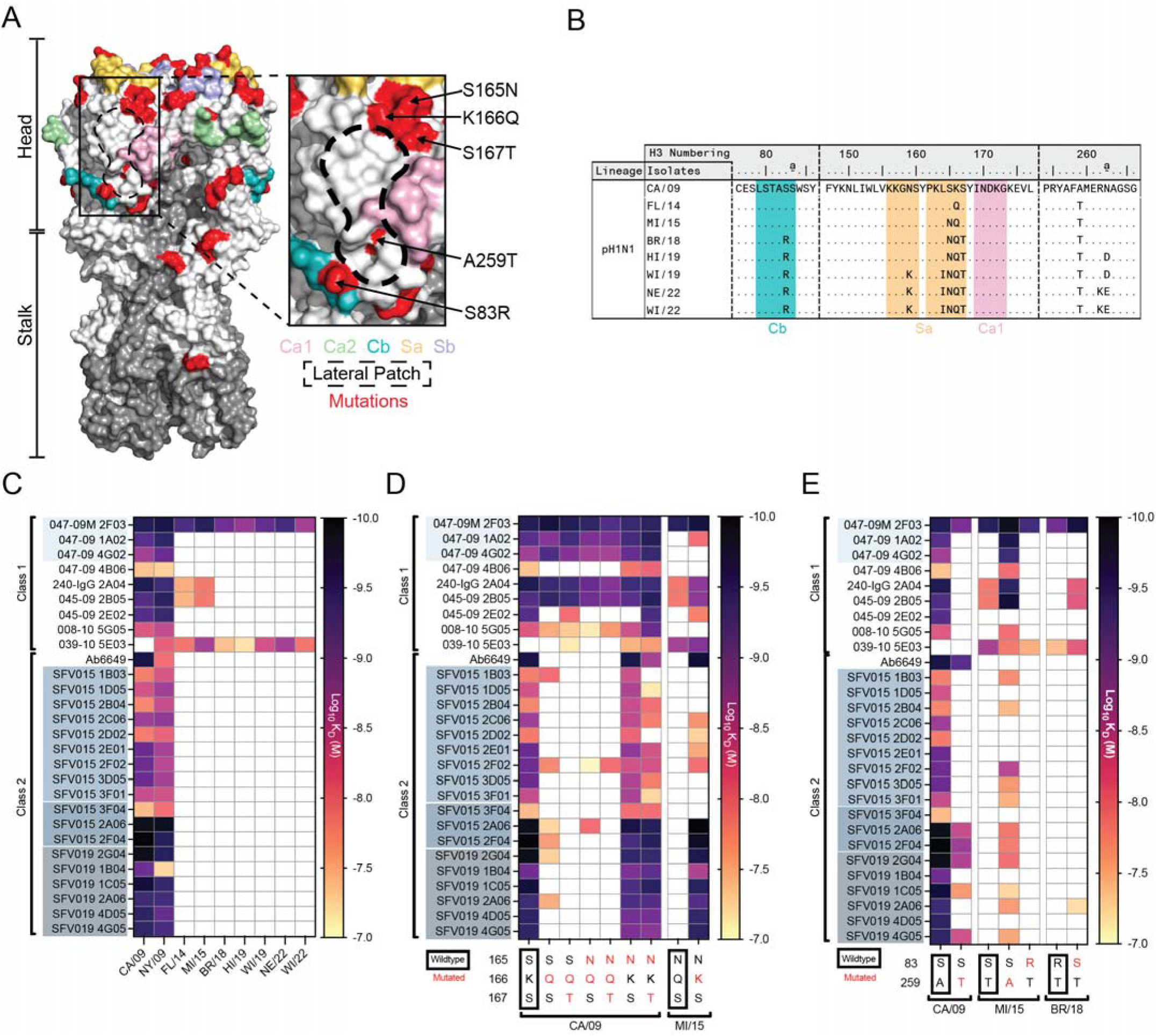
Antigenic structure and binding breadth of mAbs against the lateral patch of post-2009 H1N1 viruses. (A) Structure of HA from A/California/04/2009 (PDB: 4JTV), highlighting the major antigenic sites (Ca1, Ca2, Cb, Sa, Sb), the lateral patch epitope, and mutations in post-2009 pH1N1 viruses. The HA1 and HA2 are visualized in white and gray, respectively. (B) Amino acid sequence alignment of post-2009 H1 isolates of residues within or near the lateral patch. (C-E) Apparent affinity of lateral patch-targeting mAbs against post-2009 rH1 isolates. MAbs are divided by their class. Clonally related mAbs are highlighted by distinctly shaded blue boxes. (D-E) rH1 isolates with different mutations at positions 165/166/167 (D) and at 83 and 259 (E). White boxes represent no binding

To understand how antigenic drift in post-2009 H1N1 viruses has affected the antigenicity of the lateral patch, we evaluated the binding breadth and affinity of lateral patch-targeting mAbs against recombinant HAs (rHAs) from post-2009 H1N1 viruses (Fig. S1). MAbs used in this study were largely generated from B cells isolated from a human cohort following the 2009 pH1N1 monovalent influenza vaccine or the 2014-2015 quadrivalent vaccine (Table S3) (53,54). We also analyzed the antigenicity of the lateral patch using Ab6649, the first characterized mAb targeting the lateral patch (14). Lateral patch-targeting mAbs utilize highly restricted variable heavy (V_H_) and variable kappa (V_K_) genes (12). These mAbs can be grouped into two classes based on their V_H_ genes (Table S3) and their angle of approach based on negative stain electron microscopy (12,14). Class 1 mAbs utilized V_H_3-23, commonly paired with V_K_1-33. In contrast, Class 2 mAbs largely used V_H_ 1-2, V_H_ 3-74, or V_H_ 3-7 paired with V_K_1-33.

We found that both class 1 and class 2 antibodies lost binding to pH1N1 strains around 2014, suggesting significant antigenic drift affected mAb binding (Fig. 2C). In 2012, pH1N1 viruses acquired K166Q, located in the Sa antigenic site (Fig. 2A), which became dominant in the 2013-2014 influenza season and has persisted in circulating pH1N1 viruses (Fig. 2B).

Moreover, two additional mutations occurred directly before (S165N) and after (S167T) the K166Q mutation (Fig. 2B). These mutations occur in A/Michigan/45/2015 (MI/15; S165N only) and A/Brisbane/02/2018 (BR/18; S165N and S167T). To dissect the individual roles of these three mutations on mAb binding, we generated rHAs with various combinations of mutations at positions 165-167. We found K166Q alone dramatically reduced binding of class 2 mAbs, whereas S165N and S167T had little to no effect on binding in mutants on the A/California/04/2009 (CA/09) background (Fig. 2D). The crystal structure of mAb Ab6649 in complex with a 2006 H1 revealed that this antibody directly binds to K166, and classified as Class 2 despite containing different antibody gene repertoire (Table S3). The class 2 mAb 6649 was likely susceptible to antigenic drift at this site (14). This finding, as well as evidence that the class 2 mAb binding footprint overlaps with K166 (12), could provide an explanation as to why class 2 mAbs are more susceptible to the K166Q mutation. Reversion to Q166K in the MI/15 background restored binding by most class 2 lateral patch antibodies, and surprisingly, some class 1 mAbs.

A259T co-circulated in viruses possessing K166Q and is found near the lateral patch epitope (Fig. 2A). A259T is ubiquitous in circulating pH1N1 viruses and emerged around 2012 (Fig. 2B, Table S2). We introduced this substitution in CA/09 and found the majority of class 1 mAbs and a subset of class 2 mAbs lost binding (Fig. 2E). Reversion to T259A in MI/15 led to the restoration of binding by most class 1 lateral patch mAbs. Surprisingly, many class 2 mAbs regained binding to the T259A MI/15 despite the presence of the K166Q mutation, albeit with lower binding affinity than WT CA/09 (Fig. 2E). These data suggest that K166Q and A259T work in concert to evade the vast majority of lateral patch mAbs.

Lastly, S83R mutation emerged in the Cb antigenic site in post-2015 pH1N1 isolates and is present in the BR/18 pH1N1 isolate (Fig. 2E, Table S2). Reversion of this mutation in the BR/18 background or substitution in MI/15 revealed this mutation was necessary and sufficient for evasion of 240-IgG 2A04 and 045-09 2B05 in BR/18 (Fig. 2E). Together, these data support that distinct mutations in and around the lateral patch have affected the binding of lateral patch-specific mAbs to post-2009 pH1N1 viruses.

### Structural Analysis of Class 1 Lateral Patch-Specific mAbs

To further understand how class 1 mAbs bind to pH1N1, we determined the cryo-EM structures of mAb 047-09 1A02 (Fig S2: Table S4) and 047-09M 2F03 (Fig S3; Table S4) in complex with CA/09 HA with stabilizing mutation E47K in HA2. We compared these structures to the previously characterized 045-09 2B05 (PDB: 7MEM) (12). 047-09 1A02 and 047-09M 2F03 are clonally related, but 047-09M 2F03 exhibits improved binding breadth relative to 047-09 1A02 (Fig. 2C). 2D class averages of the complex exhibited diverse views with visible secondary structures (Fig. S2A and S3A) and three Fabs bound per HA trimer (Fig. S2B and S3B). Iterative classification and refinement in 3D (Fig. S2C and S3C; Table S4) resulted in a final map for 047-09 1A02 and 047-09M 2F03 at 3.7 Å resolution and 3.06 Å resolution, respectively (Fig. S2D and S3D; Table S4). The highest resolution features corresponded to the paratope:epitope interface and the interior of the HA head (Fig. S2E and S3E).

All three mAbs analyzed use the same heavy and light chain (V_H_3-23/V_K_1-33) and possess a Y-x-R motif within the HCDR3 that is highly conserved across class 1 lateral patch-binding mAbs (12). Despite their genetic similarities, all three mAbs exhibited distinct binding profiles. In the context of CA/09, all three mAbs made extensive interactions with the 170-loop of major antigenic site Ca1, the 80-loop of major antigenic site Cb, and residues within the 120 b-strand of the lateral patch epitope (Fig. 3A-C). Contact residues defined for each class 1 mAb share HA-antibody contact sites, specifically HA residues E119, R120, and E122 (Fig. 3A-3C). These contact residues were mapped onto a sequence alignment of post-2009 to show broadly similar HA contact sites, where the majority of these antibody contact sites fall within the heavy chain (Fig. 3A-3E). When we aligned the light chains for all three mAbs, we noted similar binding contacts except for an insertion in 047-09M 2F03. Analysis of how these mAbs orient to the CA/09 HA head revealed all three mAbs have very similar CDR loop positioning, except for the KCDR2 of 047-09M 2F03 (Fig. 3F and 3G). The KCDR2 of all three mAbs bind to HA at slightly different positions. 047-09M 2F03 KCDR2 binds to S125c via L52b, whereas 047-09 1A02 contacts S125c via E55 and Y49, and 045-09 2B05 interacts with S125C via E55. All three mAbs use the Y-x-R HCDR3 motif to bind to the epitope, with position Y100b in all three mAbs making contact with N170 (Fig. 3A-C).

**Figure 3.**
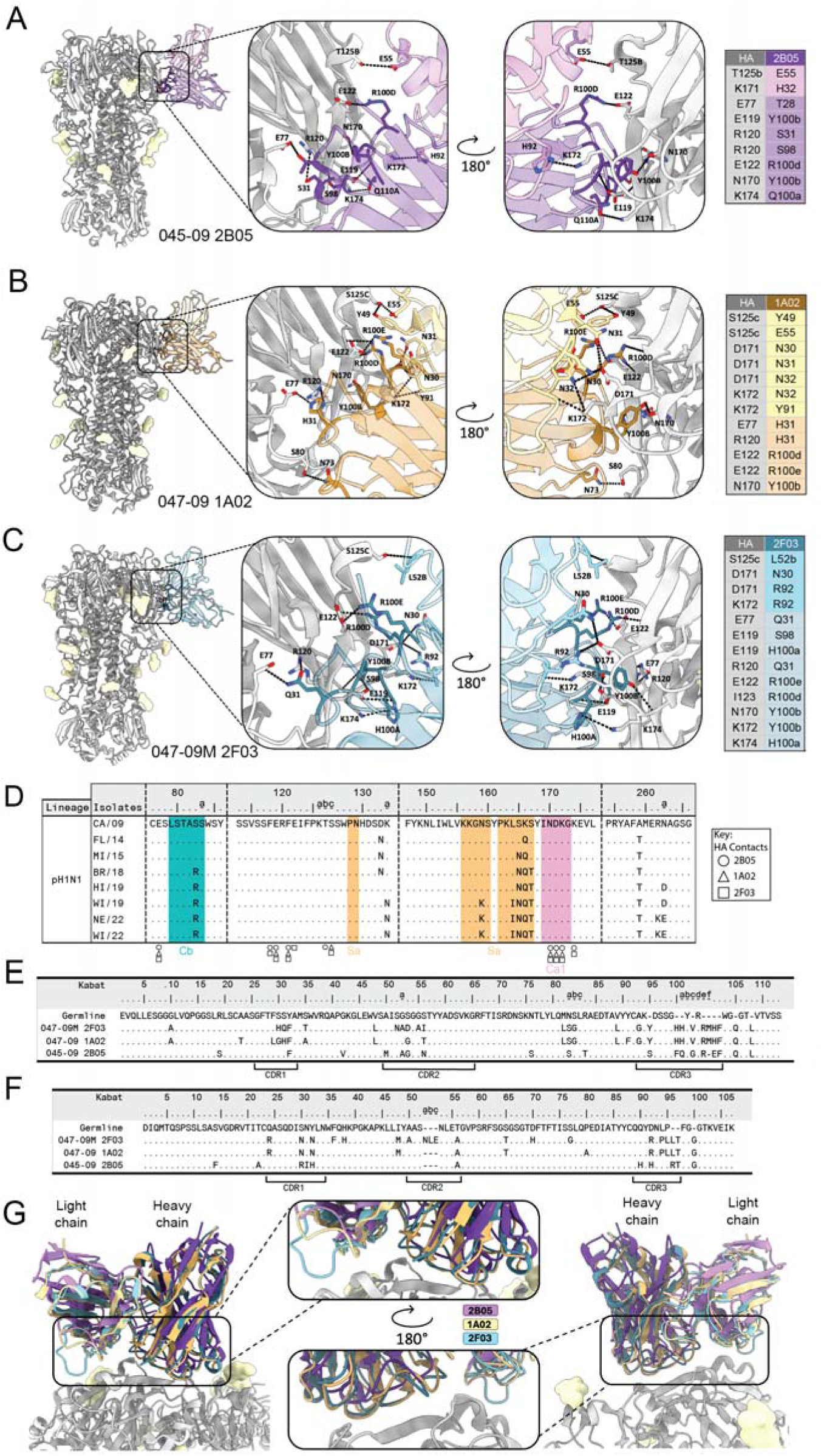
Structures of class 1 lateral patch-specific mAbs. (A-C) Cryo-EM structures of 047-09 1A02 (A), 045-09 2B05 (B), and 047-09M 2F03 (**C**) in complex with A/California/04/2009 HA. Zoomed in are the epitope:paratope interactions. Dashed lines represent contacts between the mAbs and HA. (**D**) Sequence alignment of post-2009 H1N1 viruses with mAb contacts. (**E-F**) Germline sequence alignment of 047-09 1A02, 045-09 2B05, and 047-09M 2F03 by heavy chain (**E**) and light (**F**) chains. (**G**) Overlay of the three class 1 mAbs binding to the HA head in side-to-side view and zoomed in view of the paratope.

*In vitro* viral evolution experiments revealed that 047-09 4G02, a clonal relative of 047-09 1A02 and 047-09M 2F03, drove the antibody evasion mutation A259T (12). However, the structural characterization of class 1 lateral patch mAbs revealed that A259 is neither a direct contact of this antibody nor is it surface-exposed. Rather, A259 lies on a b-strand under the 170-loop of Ca1, and this b-strand forms a b-sheet with the neighboring 120 b-strand of the lateral patch. To understand the impact of A259T and its relative position to the lateral patch β-sheets and the Ca1 loop, we performed molecular dynamics simulations of CA/09 with A259 or T259 (Fig. S4A and S4B). Introduction of A259T increased the flexibility of the 170-loop, as well as the β-strand where residue 259 resides (Fig. S4A and S4B). This predicted increase in flexibility in the 170-loop is likely due to the introduction of the bulkier and polar threonine (T) residue. The increased flexibility predicted at K174 is likely the reason we observe a decrease in binding affinity for class 1 mAbs (Fig. 2B). However, K174 is a contact for mAbs 045-09 2B05 and 047-09M 2F03 but is not a contact site for 047-09 1A02, suggesting that the A259T and the flexibility introduced at K174 may only affect certain mAbs.

### Class 2 Lateral Patch-Specific mAbs Retain Binding to Pre-2009 H1 HA Isolates

Next, we tested the conservation and antigenicity of the lateral patch in pre-2009 H1N1 viruses. We observed multiple mutations within and around the lateral patch epitope in sH1N1 viruses collected from 1933 to 2006 (Fig. 4A-B). Class 1 and 2 lateral patch-targeting mAbs cross-reacted with SC/18 and had stronger overall binding affinities than for CA/09 (Fig. 4C). Cross-reactivity to SC/18 was not surprising given that both pandemic isolates are genetically and antigenically related (12,55–57). However, these antibodies were isolated from donors born in the 1960s (Class 2) or 1980s (Class 1); their first exposure to H1 would have been an antigenically drifted sH1N1 virus. We did observe reduced binding affinity of lateral patch-specific mAbs to A/Wilson Smith/1933 (WS/33), the first human influenza isolate collected (58). This virus has been serially passaged through animal hosts and embryonated chicken eggs (59) prior to sequencing. As many mutations in WS/33 are absent from subsequent drifted variants (Fig. 4B), changes in antigenicity may be due to this serial passaging rather than antigenic drift (44,58,59). WS/33 has two mutations, S165T and K166N, relative to SC/18 (Fig. 4B), which could impact class 2 mAb binding similar to the K166Q mutation in post-2009 pH1N1 viruses. Consistent with this, WS/33 modestly reduced class 2 mAb binding (Fig. 4C).

**Figure 4.**
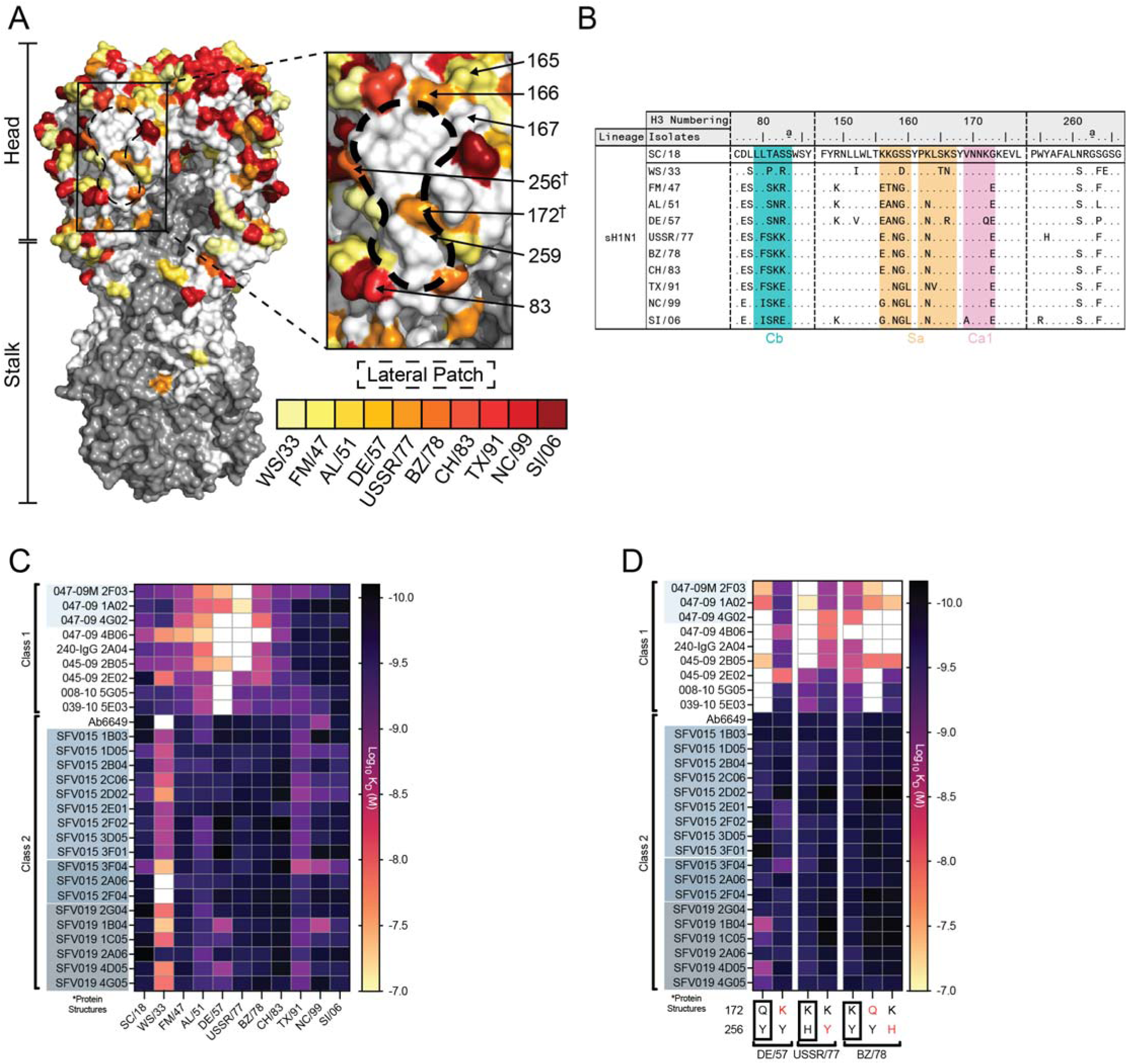
Antigenic structure and binding breadth of mAbs against the lateral patch of pre-2009 sH1N1 viruses. (A) Structure of HA from A/South Carolina/1/1918 (PDB: 1RD8), highlighting the lateral patch epitope, and mutations in sH1N1 viruses from 1918-2006. The HA1 and HA2 are visualized in white and gray, respectively. (B) Amino acid sequence alignment of pre-2009 H1 isolates, focused on residues near or within the lateral patch epitope. Colored shading depicts antigenic sites near the lateral patch. (C-D) Apparent affinity of the lateral patch-targeting mAbs screened against pre-2009 sH1N1 isolates (C) and H1 antigens with mutations at positions 172 and 256. MAbs are divided by their class. Clonally related mAbs are highlighted by distinctly shaded blue boxes. White boxes represent no binding.

Class 1 mAb binding affinity was reduced or completely ablated against influenza isolates collected from 1947 to 1957 and in 1977. Most class 1 mAbs are unable to bind DE/57 and USSR/77, notably including 047-09M 2F03, which possesses significant breadth against post-2009 viral isolates. Mutation S83R was a recurrent substitution that appeared in four isolates from 1933 to 1957 (Fig. 4D), and this mutation was reacquired in post-2015 pH1N1 isolates (Fig. 2B). Class 1 mAbs bound WS/33 and A/Fort Monmouth/1/1947 (FM/47) despite the presence of the S83R mutation. However, additional mutations arose in the Cb antigenic site, particularly A82K in FM/47, and K82N in both the 1950s isolates, A/Albany/12/1951 (AL/57) and DE/57. This mutation resulted in a predicted N-glycosylation motif site at position N82. However, position N82 was not maintained in the USSR/77 isolate since it reverted to a lysine (K) (Fig. 4D). Class 1 mAbs had reduced binding to AL/51, compared to FM/47, potentially due to the predicted N-glycosylation site introduced at K82N within the Cb epitope (Fig. 4C). Moreover, the upper portion of the lateral patch also acquired a predicted N-glycosylation motif site at residue 163 in AL/51 (Fig. 4B). Despite this, class 2 mAbs maintained binding to DE/57, suggesting the mutation at position 163, found ubiquitously in contemporary sH1N1, does not affect class 2 mAb binding (Fig. 4C). Notably, N163 is not a contact of class 2 lateral patch mAbs, suggesting why this mutation does not impact mAb binding (14). Class 2 lateral patch mAbs were able to maintain binding to all isolates, including DE/57, despite the mutation K166R (Fig. 4A and 4C).

Both DE/57 and USSR/77 isolates evaded most class 1 lateral patch mAbs. DE/57 acquired two isolate-specific mutations, K172Q at antigenic site Ca1 and K166R in Sa (Fig. 4B). In contrast, USSR/77 acquired a mutation bordering the lateral patch, Y256H, and a mutation at R83K in the Cb antigenic site (Fig. 4B). However, Q172 in DE/57 was not maintained in USSR/77, and H256 in USSR/77 was not retained in A/Brazil/11/1978 (BZ/78). We reverted the mutations at 172 and 256 in DE/57 and USSR/77, respectively. We then introduced the isolate-specific mutations K172Q and Y256H into BZ/78. Finally, we screened against lateral patch-specific mAbs to test their binding affinity for rHA. Both mutations ablated class 1 mAb binding in BZ/78, whereas DE/57+Q172K and USSR/77+H256Y restored class 1 mAb binding (Fig. 4D).

All lateral patch-targeting mAbs bound isolates from 1978 - 2006, potentially due to no mutations in the lateral patch during this timeframe that would reduce or eliminate antibody binding (Fig. 4A-D). We noted that position A259 was conserved across all pre-2009 sH1N1 isolates used in this study. Overall, this demonstrates that both classes of lateral patch-targeting antibodies retained binding to H1 antigens from 1983-2006, suggesting the lateral patch remained conserved during this period.

The two classes of lateral patch-specific antibodies are from donors with discrete birth years. Class 1 mAbs were from donors born in the mid-1980s, and class 2 mAbs from donors born in 1961 and 1975 (12,14) (Table S3). Donors with class 1 antibodies were likely imprinted with sH1N1 viruses from the late 1980s to the early 1990s, which are similar to A/Texas/36/1991 (TX/91). In contrast, donors with class 2 antibodies likely generated their primary responses against the reemerging H1N1 virus in 1977 (12,14,20). Thus, the first H1N1 virus infection may have shaped which mAb class was induced to bind to the lateral patch.

### Lateral Patch-Targeting mAbs Retain Binding to Zoonotic H1

To elucidate the conservation and antigenicity of the lateral patch of zoonotic H1 HAs, which pose a risk to human health via potential spillover events, we tested the binding breadth of lateral patch-targeting mAbs to H1 antigens of zoonotic origin. These included rHAs from 1A, 1B, and 1C swine lineages of H1N*x* viruses and avian-origin lineage H1.

We observed that both class 1 and 2 mAbs bound to 1A classical swine lineage viruses from 1931-2001 (Fig. 5A), suggesting that this epitope was antigenically conserved during this time period in swine influenza viruses (45,46). The only exception was the OH/15v isolate, which had reduced binding or completely evaded a subset of class 1 and 2 lateral patch-targeting mAbs (Fig. 5A). The OH/15v isolate was obtained from a lethal human infection with a variant 1A swine H1N1 virus (42). The OH/15v has multiple substitutions in and around the lateral patch, including I166 (Fig. 5B). As K166 is a known contact of the class 2 lateral patch mAb Ab6649 (14), it is plausible that this substitution could be responsible for reduced binding by this class of antibodies. Furthermore, the substitution R172 in the Ca1 antigenic site of OH/15v may have contributed to reduced binding by class 1 mAbs (Fig. 5B). Together, these data support that 1A classical swine viruses have maintained conservation within the lateral patch. However, the evasion of lateral patch-targeting mAbs (Fig. 5A) and ferret anti-sera (42) by OH/15v warrants further investigation into whether circulating 1A viruses have undergone antigenic drift that efficiently evades lateral patch-targeting antibodies.

**Figure 5.**
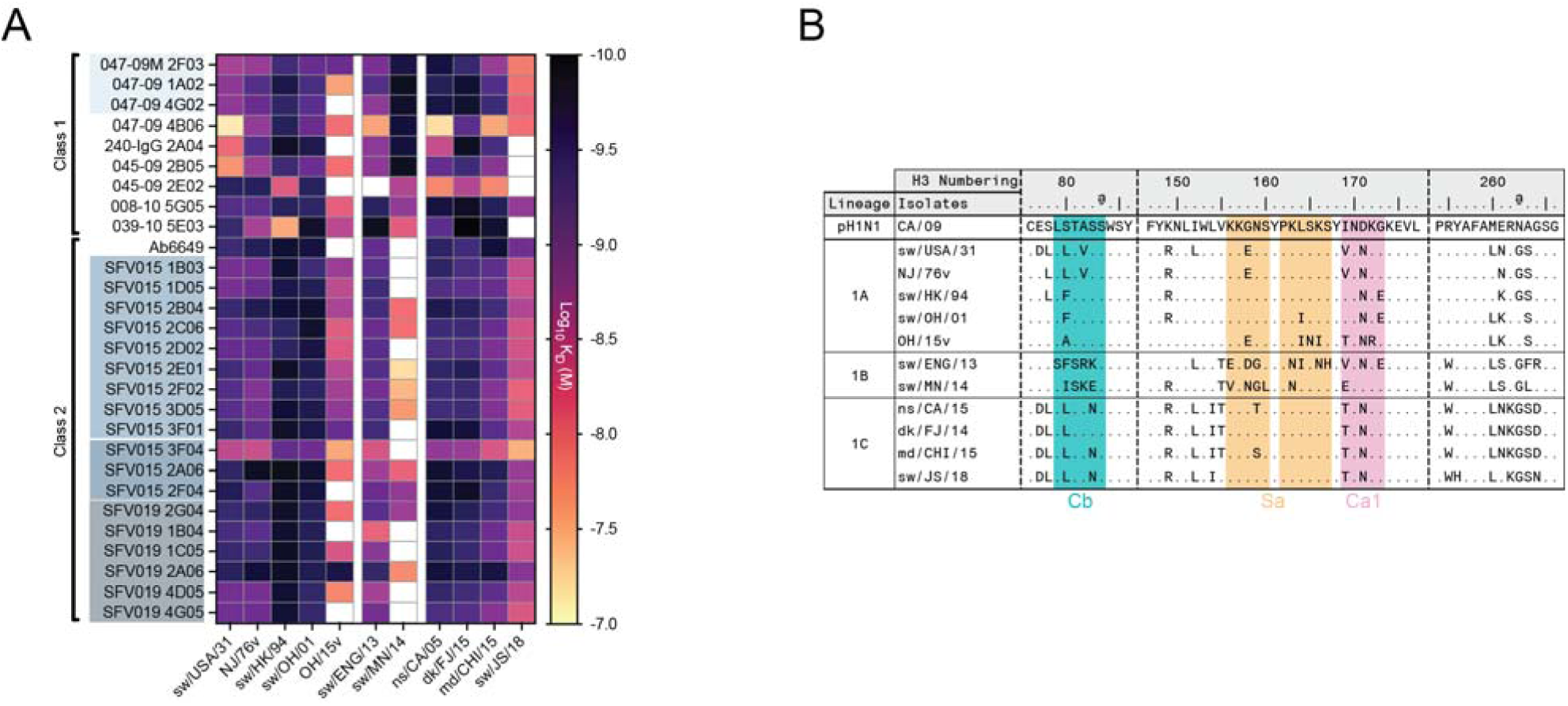
Binding breadth of the lateral patch-targeting mAbs in zoonotic H1 isolates. (A) Apparent affinity of the lateral patch-targeting mAbs screened against zoonotic H1N*x* isolates presented in a heat map. MAbs are divided by their class. Clonally related mAbs are highlighted by distinctly shaded blue boxes. White boxes represent no binding. rHA are grouped by lineages and within chronological order. (B) Amino acid sequence alignment of selected zoonotic H1 isolates focused on residues near or within the lateral patch epitope to reference strain CA/09, corresponding to mutations in positions 83, 165-167, and 259. Colored shading depicts antigenic sites near the lateral patch.

Next, we tested binding of lateral patch-targeting mAbs to two lineage 1B swine isolates. These swine isolates share common ancestors with sH1N1 derived from the 1980s and 1990s. The A/swine/England/063782/2013 (sw/ENG/13) virus is phylogenetically related to A/Chile/1/1983 (CH/83), and A/swine/Minnesota/A01567011/2014 (sw/MN/14) is phylogenetically related to TX/91 (Fig. 1B). Class 2 mAbs maintained binding to sw/ENG/13 but had reduced binding to sw/MN/14 (Fig. 5A). In contrast, class 1 mAbs bound to both antigens nearly identically. Both 1B swine isolates differ in mutations at or near the lateral patch (Fig. 5B). The lack of binding by class 2 mAbs may be due to the substitution E169 in antigenic site Ca1 (Fig. 5B), as this is a binding contact for Ab6649 (14). Although these 1B swine isolates derive from human sH1N1, their genetic and antigenic differences may be the result of divergent antigenic drift in the US and Europe (34,47,60).

Lastly, the lateral patch epitope is well-conserved in zoonotic isolates of lineage 1C, as all mAbs exhibit a high binding affinity to the lateral patch, especially to H1s of avian origin. However, about half of the class 1mAbs and several class 2 mAbs exhibited reduced binding affinity against the A/swine/Jiangsu/J004/2018 (sw/JS/18) isolate (Fig. 5A). Sw/JS/18 has a Y256H mutation, which is also found in the human isolate USSR/77 (Fig. 5B). Class 1 mAb 047-09M 2F03 was able to bind to this swine isolate, despite not binding to USSR/77 (Fig. 4C). In contrast, lateral patch mAbs 039-10 5E03 and 045-09 2E02 were able to bind to the USSR/77 but were unable to bind to sw/JS/18 (Fig. 4C and 5A). Together, these data support that the lateral patch is well conserved across zoonotic H1 viruses, but further antigenic characterization of zoonotic H1Nx viruses is necessary to determine whether these viruses are undergoing antigenic drift to evade broadly neutralizing antibodies.

### Antigenic Evolution of the Lateral Patch

Using binding data from our rHA-antibody ELISAs, we assessed the antigenic relatedness of H1 isolates by generating antigenic cartography maps (Fig. 6). Classically, these maps are generated from hemagglutination inhibition assays with human or ferret sera to visualize antigenic drift over time (49,61–63). Here, we adapted this approach using lateral patch mAb binding affinities to understand lateral patch antigenicity over time and species.

**Figure 6.**
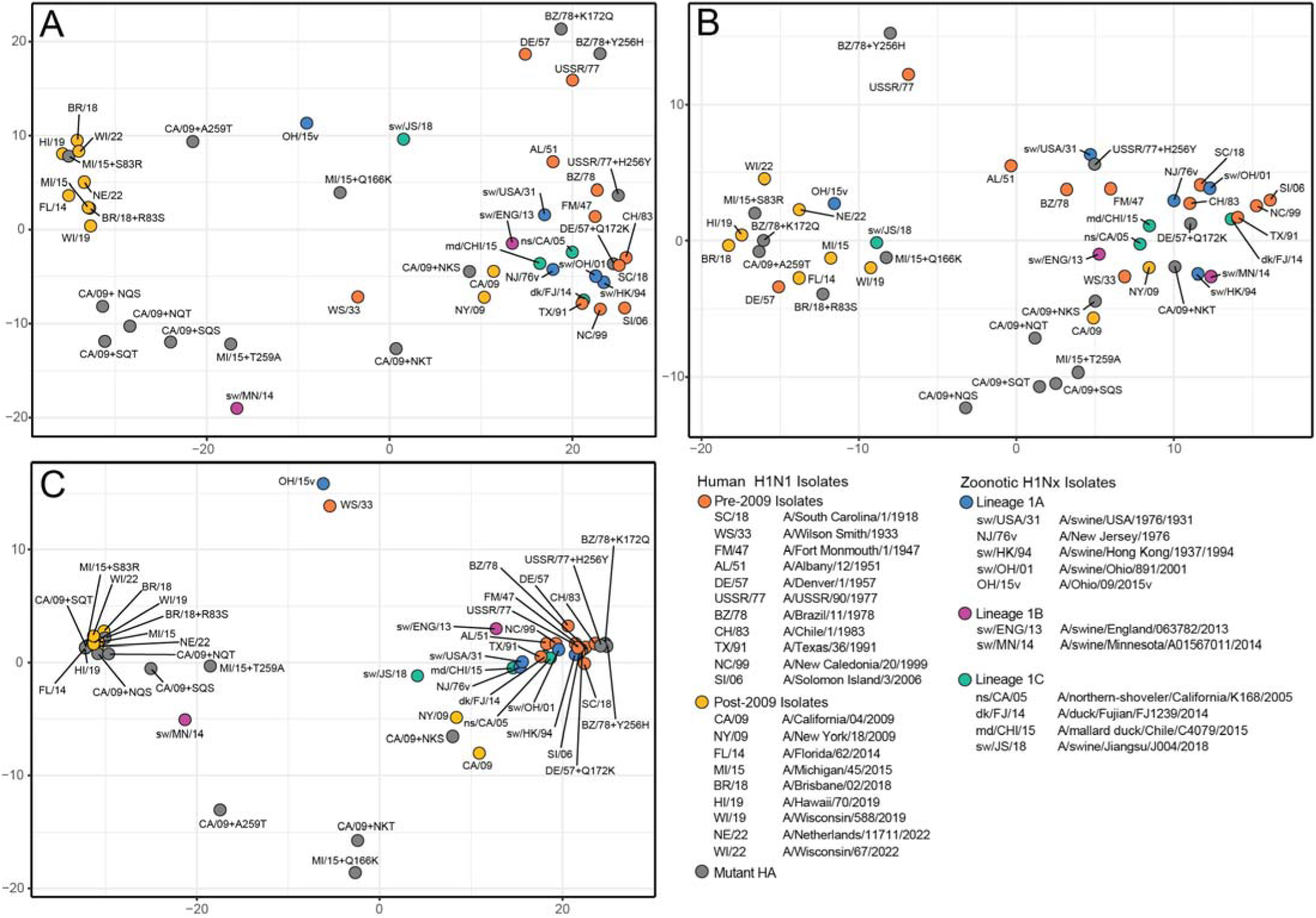
Antigenic relationship of human, zoonotic, and mutant rH1s based on lateral patch-specific mAb binding. (A) Antigenic cartography of all lateral patch-targeting mAbs against all H1 antigens. (B) Antigenic cartography of Class 1 lateral patch-targeting mAbs. (C) Antigenic cartography of class 2 lateral patch-targeting mAbs.

Analysis of an antigenic map of all lateral patch-targeting mAbs revealed that clustering patterns are largely driven by post-2009 H1N1 strains with K166Q, A259T, and S83R mutations. This cluster includes mutated post-2009 HAs that possess these mutations (Fig. 6A-B). Furthermore, three distinct sH1N1 isolates (WS/33/ DE/57, USSR/77) and the three recently collected swine isolates (sw/OH/15v, sw/MN/14, sw/JS/18) are in a unique antigenic cluster away from sH1N1, mutant sH1N1, and zoonotic isolates due to differences in mAb binding affinity (Fig. 6A).

Next, we generated antigenic maps specifically for class 1 and class 2 mAbs (Fig. 6B and 6C). HA clustering for class 1 mAbs is largely driven by the presence of A259T and S83R in post-2014 isolates, and K172Q and Y256H in sH1N1 viruses (Fig. 6B). An antigenic map of Class 2 mAbs revealed tight clusters associated with pre- and post-2009 H1N1 viruses, largely driven by the presence of K166Q. Furthermore, WS/33 harbors mutations at the Sa antigenic sites that likely drive evasion of class 2 lateral patch-targeting mAbs, leading to its separation from other pre-2009 sH1N1 viruses in this map (Fig. 6C).

Lastly, the zoonotic isolates demonstrate distinct clustering in all antigenic maps. This likely is reflective of their phylogenetic relatedness. For example, 1A classical swine and the 2009 HA isolates cluster closely with SC/18, and 1B swine viruses cluster near the 1978-2006 sH1N1 antigens (Fig. 1; 6A-C). However, swine isolates such as OH/15v, sw/MN/14, and sw/JS/18 harbor mutations in and around the lateral patch epitope that largely diminish binding to class 1 and class 2 lateral patch-targeting mAbs (Fig. 5A-5B; 6A–6C). Although K166Q in pH1N1 drives the evasion of class 2 antibodies, other substitutions found in Ca1 (E169 and R172), Sa (I166), and Y256H likely contribute to differences in antigenicity relative to human isolates. Overall, these data support that lateral patch antigenicity is distinct between post-2014 H1N1 viruses and all other human H1N1 viruses, suggesting that the induction of lateral patch antibodies by first exposure to the 2009 pH1N1 likely drove rapid and dramatic antigenic drift in this epitope. However, distinct antigenic drift events appear to have occurred in pre-1977 sH1N1 viruses and zoonotic H1s, as diversity in the zoonotic HA head domain can be highly dynamic and evolving independent trajectories across divergent swine lineages.

## Discussion

In this study, we utilized lateral patch-specific mAbs from humans vaccinated against the 2009 pH1N1 to assess their binding affinity to HA H1 strains from human isolates collected between 1918 and 2022 and zoonotic H1N*x* isolates collected from different hosts, lineages, and geographical regions. Most lateral patch-targeting mAbs in our panel bound to human H1 isolates prior to 2014, despite reduced binding affinity against isolates from 1951 to 1977. Moreover, we found that most lateral patch-targeting mAbs bound to zoonotic H1s, suggesting that this epitope is well-conserved across zoonotic viruses and could mediate protection in the event of an H1 pandemic.

Class 2 mAb Ab6649 targets the upper portion of the lateral patch, which overlaps with the Sa antigenic site (12,14). This footprint includes position K166Q, a mutation known to be universally present in post-2014 pH1N1 isolates (12,14,16). A recent study revealed that substitutions at position 128, 166, and 171 in pre-2009 sH1N1 can evade Ab6649 and its clonally related member, Ab9207 (64). Using deep mutational scanning, the authors found that Ab6649 and Ab9207 bound to a USSR/77 escape mutant possessing substitutions K166L/A. Both of these amino acids have chemical properties similar to isoleucine (I), found in OH/15v. However, these mAbs had reduced avidity in drifted strains of A/Siena/10/1989 (SN/89) and ablated binding in the further drifted strain, SI/06. These included mutations at position 166 (L/A), as well as additional substitutions at residues 126, 128, 169, 171, and 173. These positions, except for 126 and 128, are also binding contact sites for Ab6649 (14). Position 173 introduced a potential N-glycosylation site at N171, eliminating binding avidity to both mAbs 6649 and 9207 (64).

Class 1 lateral patch targeting mAbs target the lateral patch from a perpendicular angle, likely preserving binding to K166Q mutant viruses. Class 1 lateral patch mAbs have a Y-x-R motif within the HCDR3 that makes multiple critical contacts with the lateral patch. A prior study revealed that class 1 lateral patch mAbs can drive mutations at positions E119, R120, and A259, with the prior two mutations being contacts of the Y-x-R motif (12). A recent study of three clonally related class 1 lateral patch mAbs revealed the mAbs could drive escape mutations at positions E119, R125, and K172, as well as the introduction of N-glycosylation motifs at positions P122, K123, and E124 (65). All of these sites are critical contacts of the Y-x-R motif in class 1 mAbs utilized in our study and the Y-x-K motif in their selected mAbs (12,65). Notably, residue P122 emerged as an important contact residue in that study, in which viral escape occurred readily. Moreover, it is an important contact of 045-09 2B05, a class 1 lateral patch antibody (12). Yet another class 1-like mAb, 441D6, was elicited from a mosaic receptor-binding domain of human H1 HA on a nanoparticle in mice (66). MAb 441D6 demonstrated strong neutralization potency against a panel of H1 HA strains collected over 90 years, except for A/Iowa/1943. However, mutations such as K172E and E119K significantly reduced binding (66). Despite these potential evolutionary trajectories, only A295T has emerged and persisted in circulating pH1N1 viruses. Overall, these studies highlight how diverse substitutions at different residues in the *in vitro* escape mutants of human H1 HA lateral patch and lateral patch-direct lineages can affect binding affinity and neutralization potency.

Despite the significance of how specific substitutions at different positions affect antibody binding in our data, the broadly reactive H1 lateral patch mAb 047-09M 2F03 bound to all post-2009 H1 HA isolates and many human H1 isolates since 1918. We found that 047-09M 2F03 was evaded only by Y256H found in USSR/77. Y256 is stacked between two arginine residues within the lateral patch epitope. The Y256H mutation could induce a pH-dependent conformational change, as protonation of histidine can result in repulsive interactions with the two arginine residues and thereby increase the general flexibility within the binding interface. Our data show that K172Q in DE/57 and Y256H in USSR/77 reduced class 1 mAb binding. pH1N1 clade 5a.2a has acquired a K172Q mutation, which could potentially impact class 1 lateral patch antibody binding. As K172 is a contact of class 1 lateral patch mAbs, it is possible that this mutation could be responsible for antigenic drift within the lateral patch of contemporary viruses (43,67). 047-09M 2F03 is part of the same clonal family as 047-09 4G02 and 047-09 1A02, though we observed that these latter mAbs do not bind to post-2014 H1N1 viruses due to the A259T mutation. Moreover, other class 1 lateral patch-targeting mAbs, such as 240-IgG 2A04 and 045-09 2B05, retain binding to A259T mutants, but lose binding to HA with S83R. R83 may clash with 045-09 2B05 and 240 IgG 2A04, leading to the reduced binding we observed. It remains unknown what mechanism allows 047-09M 2F03 to retain binding to post-2014 H1N1 viruses. Overall, epitope-specific mAbs are sensitive to distinct lateral patch mutations despite overlapping binding footprints and identical or very similar gene usage.

Our study supports the notion that the lateral patch was well conserved for nearly 100 years, with the exception of mutations arising in 1950s and 1977 sH1N1 viruses that evade class 1 antibodies and post-2014 pH1N1 viruses. Conserved sites in HA are often maintained due to functional constraints. However, the lateral patch lacks any described or exhibited function (14), unlike the RBS, another conserved epitope on the HA head domain. Understanding the function, or lack thereof, of the lateral patch could reveal why this region has been well conserved across human and zoonotic H1Nx isolates.

The effect of glycans may play a role in reducing antibody responses to the HA head, as the sH1N1 and pH1N1 HA have accumulated multiple N-linked glycans over time. In sH1N1, N-glycosylation sites temporally appeared at positions N129 (1986–2008), N131 (pre-1957;1977-1986), N158 (pre-1957 and 1977-1986), and N163 (pre-1957; 1977-2008); whereas in pH1N1, one characterized N-glycosylation site appeared in post-2014 pH1N1 (16,67,68). All of these glycosylation sites were located within and around the Sa antigenic site. However, our lateral patch mAbs were able to bind despite the likely presence of introduced N-linked glycans at position S165N in post-2014 pH1N1 viruses and temporally positioned glycans in 1977-2006 sH1N1. Donors primed with pre-1985 sH1N1 showed a K166-immunodominant response prior to the acquisition of the N-glycosylation site at position 129 in post-1985 sH1N1 (16). It is noted that N129 is predicted to shield K166 in the hemagglutination-inhibition assay and that this effect was reproducible in an animal study (66). Our data show that class 2 mAbs were from donors born in 1961 and 1975 and are K166-specific mAbs (12,14). However, despite the temporally positioned glycans at the Sa antigenic site, specifically N129, class 2 mAbs bound to three isolates collected between 1991 and 2006. N-glycosylation plays an essential role in evading antibody responses, as it shields antigenic sites (69,70), and IAVs commonly acquire N-linked glycans as they drift (69,71). It is unclear if these mAbs bind to N-linked glycans near the lateral patch or have affinity matured to evade these glycans. However, a study found that N165, Q166, and I219T were crucial for completing the N-glycosylation site at N165 (72). Both N165 and I219T co-occurred in the 2015-2016 season, as did K166Q and A259T during the 2013-2014 influenza season (16,72). However, the residue sits across protomers within the HA trimer, which remains inconsistent with the N-glycosylation sequencing pattern and epistatic interactions (72). It remains unknown whether I219T plays a role in N-glycosylation acquisition and whether it affects antibody binding.

The introduction of IAVs into the human population resulted from zoonotic spillover from swine and avian species (5,10,38). Currently, numerous surveillance studies have been conducted, highlighting the diverse influenza virus subtypes, clades, and lineages that circulate in distinct regions of the world (73). In addition to the spillover of zoonotic influenza viruses into humans, reverse zoonosis of human influenza viruses back into animals, particularly swine, has created reservoirs of IAVs adapted to humans. Notably, multiple reverse zoonosis events have occurred after the 2009 pH1N1 outbreak, leading to the 1A.3.3.2 lineage in swine worldwide (30,31,49,63,74,75). Multiple substitutions at position 166 in the Sa epitope have been observed in swine H1 clades (49,74,76). Furthermore, there are distinct amino acid mutations in the Ca1 and Cb antigenic sites in the Korean 1A.3.3.2 clade, subclades clade I and II, including K172R and S83K/R, respectively (75). Despite this, the rate of amino acid substitution in swine H1N*x* remained low compared to human H1N1 viruses (77–79). As antigenic drift occurs more slowly in animal reservoirs due to reduced lifespan and immune pressure on IAVs, this would explain why the lateral patch in zoonotic isolates remains conserved. However, reassortment events of H1N*x* subtypes with other diverse IAVs can increase the genetic and antigenic diversity of H1 swine lineages, independent of the lateral patch (75,80–82).

Studies using age-stratified human sera against swine H1 subtypes have revealed that individuals born in different years retain cross-reactivity against specific subclades in the 1A and 1B lineages (63,83). This suggests that specific age cohorts can have cross-reactive antibody responses in the event of a spillover. However, Eurasian avian-like H1N*x* viruses remain a concern, as there is minimal cross-reactivity to this lineage, and these viruses continue to evolve across distinct species (50,81,84–87). Swine 1C viruses, such as sw/JS/18, are known to have pandemic potential, as they have high genetic diversity, show efficient infectivity and aerosol transmission in ferret models, and there is little to no cross-reactive antibodies in human sera against these viruses (81,84).

As H1N*x* remains in circulation in swine and wild avian species, the recurrence of pandemic potential H1N*x* virus in humans remains high, as multiple cases of H1N*x* variants (H1N*x*v) have been detected and can cause severe disease (42,74,88). However, our data support that lateral patch-targeting mAbs largely retain considerable binding breadth against 1A, 1B, and 1C swine H1N*x* viruses, and memory B cells encoding these antibodies could be harnessed to protect in the event of an outbreak or pandemic. Ongoing surveillance of circulating H1N*x* viruses in zoonotic reservoirs can inform the suitability of inducing lateral patch-targeting antibodies with vaccines and the potential of lateral patch-targeting antibodies as prophylactics and therapeutics.

In summary, antigenic evolution in H1-expressing influenza viruses is dynamic, as the requirement to evade antibody responses from different species never abates. Ongoing surveillance systems for swine and wild avian species are necessary to assess how H1 has evolved to escape human immunity. Our study provides an analysis of a unique, conserved epitope—the lateral patch—which may offer broad protection in the event of a spillover. Together, screening neutralizing mAbs against structurally variable and zoonotic H1s can be used to inform and design next-generation prophylactics or therapeutics.

## Acknowledgement

We thank members of the Guthmiller and Ward labs for critical feedback on this study. We also thank Drs. Ian Wilson and Xueyong Zhu for helpful discussions on the structures of lateral patch-targeting mAbs. We thank Dr. Tavis Anderson for the discussion on swine viruses and on antigenic cartography analyses.

## Author Contributions

JJG and JLR conceptualized the study. JJG and AW acquired funding and provided resources. JJG, JLR, MG, WJ, MFQ, and JH designed and developed the experimental methodology. JLR, MG, WJ, DS, OS, MFQ, and JH conducted the project investigation. JJG and AW provided supervision of the project. JLR and JJG conducted the visualization and writing. Revision and editing were performed by JLR, MG, JJG, MFQ, and JH.

## Data availability

All data to understand the conclusions of this research are available in the main text, supplementary materials, or repositories listed below. 047-09M 2F03 and 047-09 1A02 with CA/09 model were deposited onto the Electron Microscopy Data Bank (EMD-76461 and EMD-76462, respectively) and the Protein Data Bank (PDB: 12IU and PDB: 12IV, respectively).

## Methods and Materials

### Cell Culture

Human embryo kidney (HEK) 293T cells (Thermo Fisher Scientific) were maintained at 37°C with 5% CO2 in 15-cm surface-treated tissue culture dishes (Fisher Scientific) in Advanced Dulbecco’s modified Eagle’s medium (DMEM; Gibco) supplemented with 2% ultralow IgG fetal bovine serum (FBS; Gibco), 1% L-glutamine (Gibco), and 1% antibiotic-antimycotic (Gibco). These conditions were maintained throughout the protocol by splitting at approximately 80% confluency.

Expi293F cells (Thermo Fisher Scientific) were maintained at 37°C, 125 rpm, 8% CO2 in advanced with 80% humidity in plastic flasks with ventilated caps (Avantor VWR® Erlenmeyer Flasks, Sterile, Polycarbonate with 0.22 μm ventilated caps). An OrbiShaker™ CO2 shaker with 19 mm orbitals (Benchmark Scientific) was used for aeration and mixing. These conditions were maintained throughout the protocol. During the maintenance and expansion phase, the cells were split to a concentration of 0.3-0.4 10^^6^ viable cells/ml when they reached a density of 3–5 × 10^^6^ viable cells/ml. This required splitting roughly every 4th day post-subculture. Cell density was determined with a Countess™ 3 FL Automated Cell Counter (Thermo Fisher Scientific).

### Protein Production

#### Monoclonal Antibody Generation

MAbs used in this study are from other published studies (12,14,53,54). Immunoglobulin heavy and light chain plasmids were generated from the previously described study (12,14). Plasmids for the heavy and light chains of a corresponding mAb, 9 μg each in 2.4 ml of DMEM (Gibco), were mixed with 100 μg of 1 mg/ml of polyethyleneimine (PEI; Sigma-Aldrich), mixed well by vortexing, and co-transfected into confluent HEK 293T cells for 12 to 18 hours, after which the media were changed to Protein-Free Hybridoma Medium (PFMH-II; Gibco). The supernatant was harvested and clarified after 5 days. The secreted mAbs were purified from the supernatant by passage over Protein A agarose beads (Pierce), followed by sequential centrifugation and washes with 1M NaCl and 1x phosphate-buffered saline (PBS). MAbs were eluted using glycine-HCl (0.1 M) and neutralized with Tris-HCl (1 M). Proteins was concentrated and buffer exchanged using Amicon Ultra Centrifugal filter (Sigma) in 1x PBS. Protein concentration was measured with Nanodrop One C Spectrophotometer (Thermo Scientific).

#### Recombinant HA

Recombinant HA amino acid sequences were obtained from Global Initiative on Sharing All Influenza Data (GISAID) and cloned in a plasmid with a fold-on trimerization domain, AviTag™, and 6xHisTag. All constructs possessed a Y98F (H3 numbering) mutation to reduce sialic binding (89). 9 μg of plasmid DNA in 2.4 ml of DMEM (Gibco), was mixed with 100 μg of 1mg/ml of PEI (Sigma-Aldrich), mixed well by vortexing, and co-transfected into confluent HEK 293T cells for 12 to 18 hours, after which the media were changed to PFHM-II (Gibco). The supernatant was harvested and clarified via centrifugation at the end of the 4^th^ day. Expi293F cells followed a different protocol to prepare a DNA-PEI complex by mixing 100 μg of DNA with 500 μl of PEI in 5 ml of Freestyle 293 Expression Medium (Gibco) (per 100 ml of cell culture) and co-transfected into high-density Expi293F cells for 12 to 18 hours, after which the media were switched with fresh Expi293 expression medium. The supernatant was harvested and clarified via centrifugation at the end of the 3^rd^ day. The secreted HAs were purified from the supernatant using Ni-NTA agarose (Qiagen), followed by gravity-flow purification with disposable 10 ml columns (Fisher). The supernatant-resin mixture was added at a controlled flow rate and washed with washing buffer (20 mM NaH_2_PO_4_, 500 mM NaCl, and 20 mM imidazole at pH 7.4) and elution buffer (20 mM NaH_2_PO_4_, 500 mM NaCl, and 500 mM imidazole at pH 7.4). The eluate is collected, buffer exchanged, and concentrated using an Amicon Ultra Centrifugal filter (Sigma) with 1X PBS. The protein concentration was measured with bicinchoninic acid (BCA) protein assays (Pierce) and stored at −80°C.

### Cloning, Transformation, and Expression of Mutant HAs

The recombinant ectodomains of H1 (rHA) were constructed using site-directed mutagenesis. The combination of mutations found in and around the lateral patch was substituted in rH1 antigens of A/California/04/2009 (CA/09) at the naturally occurring SKS→NQT motif site and 259. Then, selected mutations were introduced in positions 83, 166, and 259 in A/Michigan/45/2015 (MI/15) and 83 in A/Brisbane/02/2018 (BR/18). Furthermore, positions 172 and 256 were selected for mutation in the respective isolates of A/Denver/1/1957 (DE/57), A/USSR/90/1977 (USSR/77), and A/Brazil/11/1978 (BZ/78).

The PCR samples were transformed into NEB Alpha-competent *E. coli* cells, following the NEB instruction manual, and spread 200 μl of each dilution onto a selection plate containing carbenicillin. The plate was incubated at 37°C overnight. Followed the instructions for Qiagen Plasmid Plus 96 Miniprep Kit from picked colonies to prepare for DNA samples. Samples were measured for concentration using a NanoDrop One C Spectrophotometer prior to Sanger sequencing (QuintaraBio). The plasmid was maxiprepped using the ZymoPURE II Plasmid Maxiprep Kit, and a glycerol stock was prepared. Constructed DNA plasmids were cloned and transfected into HEK 293T or Expi293F cells, following the methods and materials as described above.

### Antigen-Specific ELISAs

Recombinant soluble ectodomain HA was coated on a 96-well high-binding microplate (Corning) at a 2 μg/ml concentration with 1x PBS overnight at 4°C. The plates were then washed with 1x PBS containing 0.05% Tween and blocked with 1x PBS supplemented with 20% FBS for 1-2 hours at 37°C. Antibodies were then serially diluted 1:3 starting at 10 μg/ml and incubated for 1-2 hours at 37°C. Horseradish peroxidase (HRP)-conjugated goat anti-human IgG antibody (diluted 1:1000; Jackson Immuno Research) was used to detect mAb binding, and plates were subsequently developed with Super Aqua Blue ELISA substrate (Invitrogen). Absorbance was measured at 405 nm on a microplate spectrophotometer (BioTek). To standardize the assays, control antibodies with known binding characteristics were included on each plate, and the plates were developed when the control absorbance reached 3.0 ± 0.1 optical density (OD) units. All experiments were performed in duplicate. Affinity measurements, as represented by the dissociation constant (Kd) at molar concentrations (M), were calculated in Prism 9 (GraphPad) using nonlinear regression analysis.

### MD Simulations

As starting structures for our simulations, the available X-ray structure of A/California/04/2009 was used (PDB accession code: 4M4Y). For the A259T mutant, this residue was mutated in MOE using the Protein Builder tool, followed by a local energy minimization. For our simulations, we capped the N- and C-terminal parts of each domain with acetylamide and N-methylamide to avoid perturbations by free charged functional groups. For each HA, we performed three repetitions of 1000 ns of classical molecular dynamics simulations using the AMBER 24 simulation software package which contains the pmemd.cuda module. The structures were prepared using CHARMM-GUI. The structure models were placed into cubic water boxes of TIP3P water molecules with a minimum wall distance to the protein of 12 Å and the charge was neutralized with K^+^ Cl^-^ ions up to a concentration of 0.15 mM. Parameters for all simulations were derived from the AMBER force field 14SB. Each system was carefully equilibrated using a multistep equilibration protocol. Bonds involving hydrogen atoms were restrained using the SHAKE algorithm, allowing a timestep of 2.0 femtoseconds. The systems’ pressure was maintained at 1 bar by applying weak coupling to an external bath using the Berendsen algorithm. The Langevin Thermostat was utilized to keep the temperature at 300K during the simulations.

### MD Analysis

For the two investigated HA variants, we calculated the residue-wise B-factor for all frames in the MD simulation, as measure of global flexibility implemented in cpptraj to identify and quantify differences in the conformational diversity between the HA variants. Representative cluster structures of the different HA variants were obtained using the average-linkage hierarchical clustering implemented in cpptraj using a Cα-RMSD distance cut-off of 2 Å. We used PyMOL and ChimeraX to visualize protein structures and electron density maps (PyMOL - The PyMOL Molecular Graphics System, Version 3.0 Schrödinger, LLC).

### Cryo-EM sample preparation

HA complexes were prepared by mixing Fab respectively with a 3:1 molar ratio of Fab:HA (i.e., 3 Fabs per HA trimer) and incubated for 1 hour at room temperature. 0.1% w/v octyl-beta-glucoside detergent was added to the complex to aid in particle tumbling. The final concentration of sample was 1 mg/mL on the grid. A Vitrobot Mark IV system was used for the preparation of cryoEM grids. The settings were as follows: temperature inside the chamber was 4°C, humidity was 100%, blotting force was 1, wait time was 5s, blotting time was varied within a 3.5 and 4s range. 3 μL of sample was added to plasma cleaned 1.2/1.3 copper Quantifoil 300 mesh grid. The plasma cleaning step was performed in the Solarus 950 plasma system (Gatan) with Ar/O2 gas mix for 25s. The sample was blotted off for 5s and the grids were plunge-frozen into liquid-nitrogen-cooled liquid ethane.

### Cryo-EM data collection, processing, and model building

Cryo grids of A/California/04/2009 with E47K stabilizing mutation in HA2 complexed with FAbs 047-09-1A02 and 047-09M 2F03 were imaged at 190,000× nominal magnification using a Falcon 4i camera on a Glacios microscope at 200 kV, respectively. Automated image collection was performed using EPU from ThermoFisher. Images were aligned, dose-weighted, and Contrast Transfer Function (CTF)-corrected in the CryoSPARC Live™ software platform, with automated image collection also performed using Smart EPU software (ThermoFisher). Data processing was carried out in CryoSPARC v4.7.1. Micrographs with CTF fits better than 10.5 Å were selected for further processing, yielding 5,616 micrographs for 047-09M 2F03 and 5,152 micrographs for 047-09-1A02. Initial blob-based particle picking was performed on the first 100 micrographs using a minimum particle diameter of 100 Å and a maximum of 200 Å. These particles were subjected to 2D classification to generate templates, which were then used for template-based picking across all remaining micrographs.

Particles were extracted using at a 512 pixels box size and subjected to additional rounds of 2D classification. High-quality 2D class averages from the template-picked particles were used as input for topaz training, enabling the selection of fully occupied HA trimer particles and the generation of an initial 3D reference via ab initio reconstruction. This initial model was further refined through heterogeneous refinement, yielding a single high-quality class that was subsequently refined using non-uniform (NU) heterogeneous refinement. Final map resolutions determined by gold-standard Fourier shell correlation (GSFSC), were 3.06 Å for 047-09M 2F03 and 3.93 Å for 047-09-1A02. Initial coordinates for the HA model were taken from the available X-ray structure (PDB accession code: 4M4Y), and the initial antibody structure models for 047-09-1A02 and 047-09M 2F03 were predicted using the tool ImmuneBuilder.

We docked the models into the cryo-EM density map in UCSF ChimeraX. The structure model was built iteratively with COOT followed by real-space refinement in PHENIX package. The Kabat numbering system (https://www.scholars.northwestern.edu/en/publications/sequences-of-immunoglobulin-chains-tabulation-and-analysis-of-ami) (90) was used for 047-09-1A02 and 047-09M 2F03 mAbs and H3 numbering scheme for HA.

### Structural Biology and Sequence Alignment

A crystal structure was obtained from the Protein Data Bank (PDB) to map out the selected mutations in the structure of the HA from the influenza A/California/04/2009 H1N1 virus (PDB: 4JTV) and A/South Carolina/1/1918 (PDB: 1RD8) with PyMOL. The mutations were mapped using the publicly available Nextstrain dataset, focusing on the HA1 H1N1pdm genotype and amino acid evolution from 2009 to 2020, with H3 numbering.

### Phylogenetic Analysis

Sequence alignments were manually constructed for the HA H1 amino acid sequences using the MEGA12 platform. The neighbor-joining method were inferred for the amino acid sequences (91). The percentage of replicate trees in which the associated taxa clustered together in the bootstrap test (1,000 replicates) are shown next to the branches (92). The tree is drawn to scale, with branch lengths in the same units as those of the evolutionary distances used to infer the phylogenetic tree. The evolutionary distances were computed using the Johns-Taylor-Thornton (JTT) matrix-based method (93) and are in units of the number of amino acid substitutions per site. The rate variation among sites was modeled with a gamma distribution (shape parameter = 5.00). The analytical procedure encompassed 31 amino acid sequences. The pairwise deletion option was applied to all ambiguous positions for each sequence pair, resulting in a final data set comprising 566 positions. Evolutionary analyses were conducted in MEGA12 (94,95) utilizing up to 7 parallel computing threads.

### Antigenic Cartography

Using antigen-specific ELISA M Kd values generated from monoclonal antibody binding to distinct HAs, a 2-dimensional (2D) antigenic map was constructed using the Racmacs R Library for antigenic cartography (https://acorg.github.io/Racmacs/articles/intro-to-antigenic-cartography.html#converting-a-distance-table-to-a-map). Kd values that were marked as “unstable,” and had a value of more than 9_E_-7 were replaced with “9_E_-7” to limit between moderate binding and no binding. Only Kd values less than 9×10^-7^ were retained.

## Supplementary Figures

**Supplementary Figure 1:**
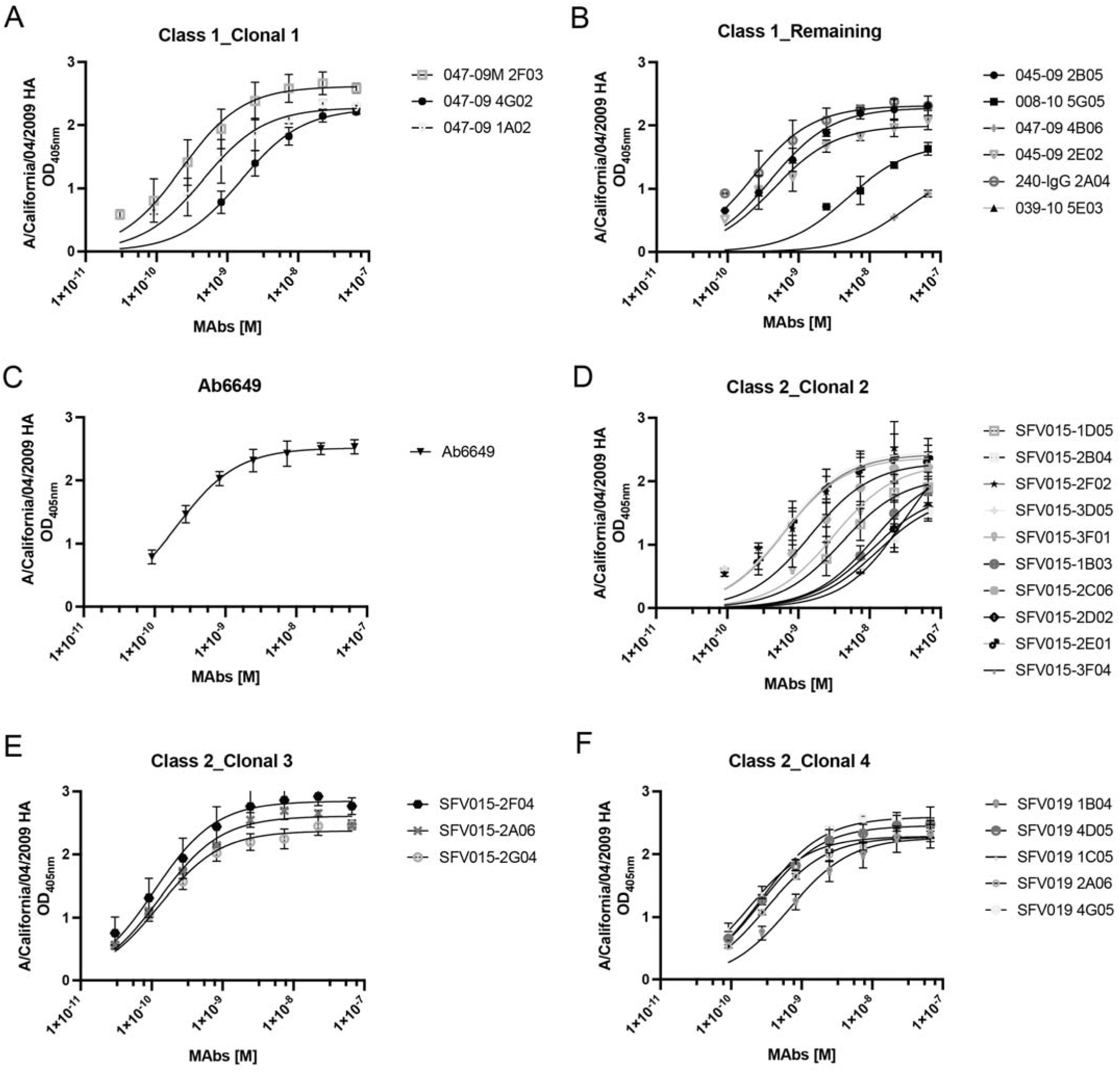
ELISA-based binding analyses of MAbs [M] screened against A/California/04/2009. Binding curves were fitted to determine the apparent binding affinity of class 1 (A-B) and class 2 (C-F) and to assess their clonality.

**Supplementary Figure 2.**
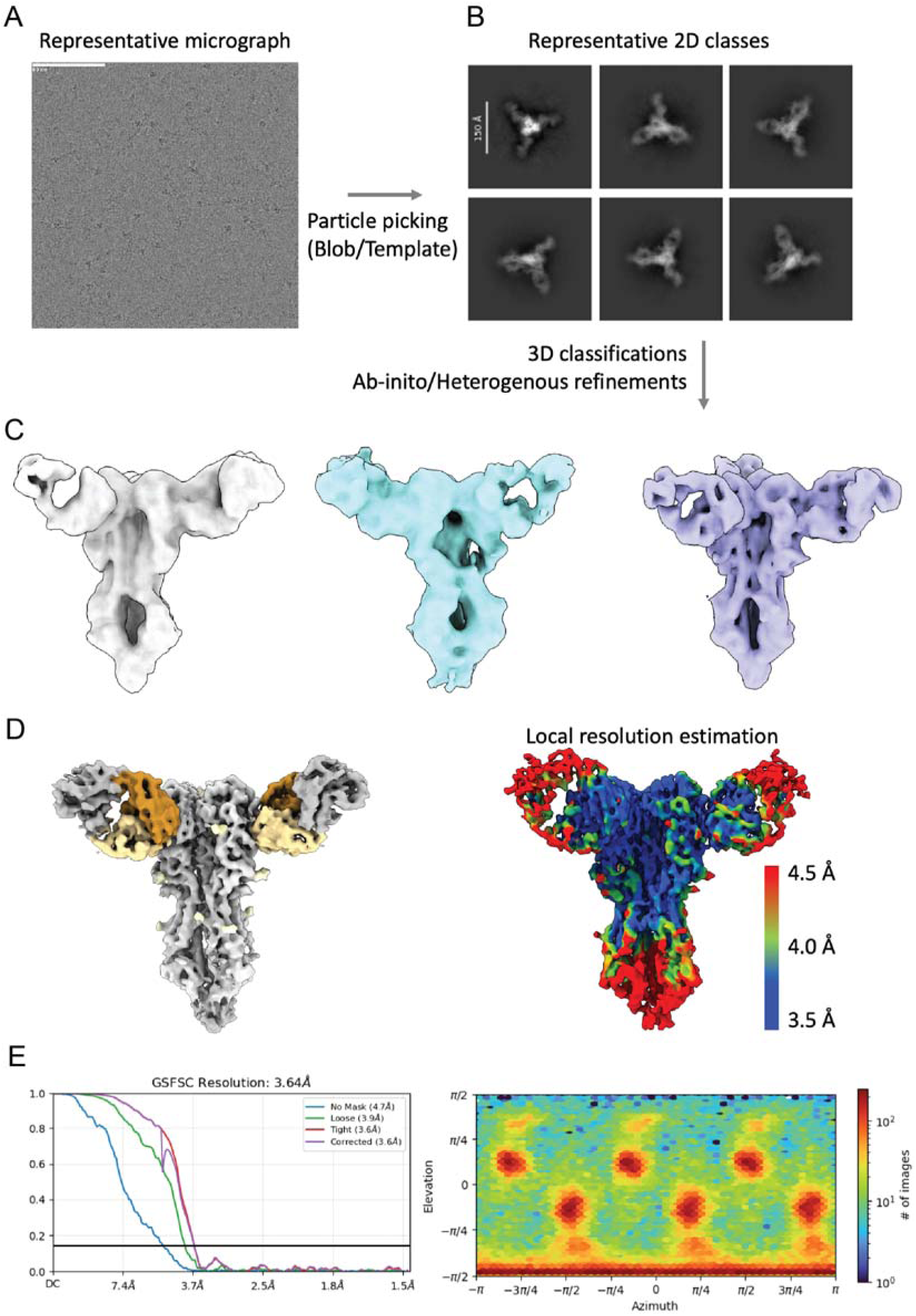
Cryo-EM data and processing and validation of 047-09 1A02 in complex with A/California/04/2009 HA. (A) Representative micrographs of mAb 047-09-1A02 in complex with A/California/04/2009 HA. Scale bar 80 nm. (B) Representative 2D class averages. Scale bar, 150 Å. (C) Ab-initio and heterogeneous refinements (representative classes shown), Non-Uniform (NU) refinement on the best class with C3 symmetry. (D) The final map is colored based on the local resolution estimation. (E) Angular distribution plot from the final NU refine job and Fourier Shell Correlation of the Electron density map.

**Supplementary Figure 3.**
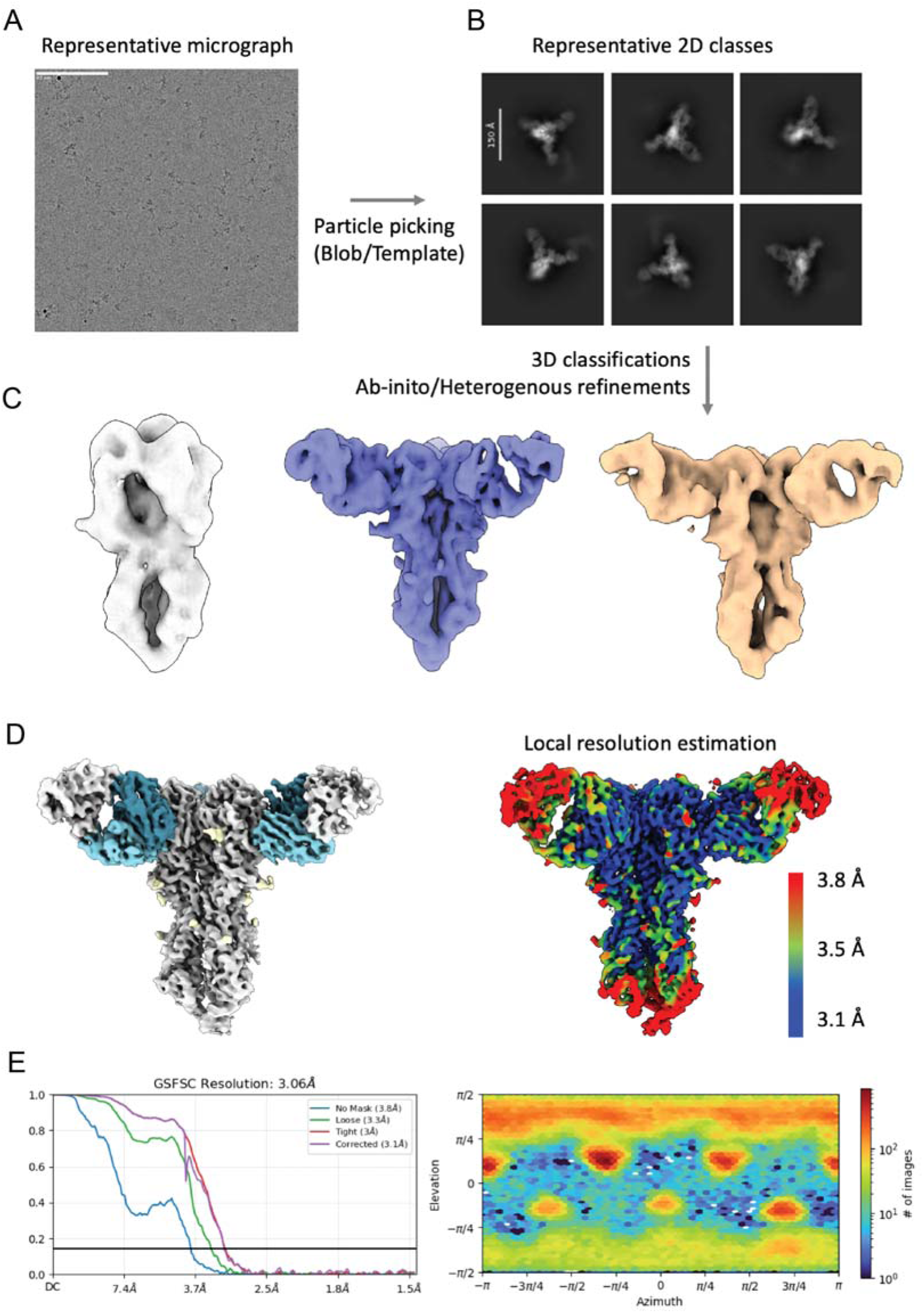
Cryo-EM data and processing and validation of 047-09M 2F03 in complex with A/California/04/2009 HA. (A) Representative micrographs of mAb 047-09M-2F03 in complex with A/California/04/2009 HA. Scale bar 80 nm. (B) Representative 2D class averages. Scale bar, 150 Å. (C) Ab-initio and heterogeneous refinements (representative classes shown), Non-Uniform (NU) refinement on the best class with C3 symmetry. (D) The final map is colored based on the local resolution estimation. (E) Angular distribution plot from the final NU refine job and Fourier Shell Correlation of the Electron density map.

**Supplementary Figure 4.**
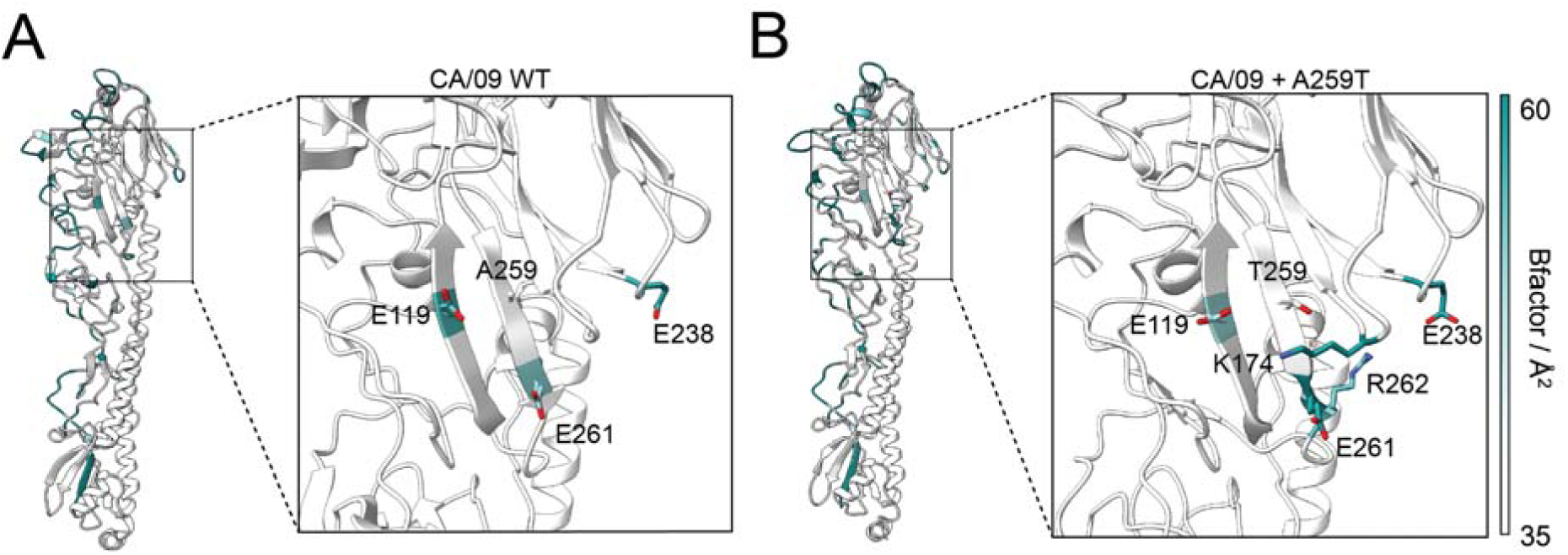
Molecular Dynamics analysis of WT HA and A259T variant. MD analysis of the flexibility of WT (A) and A259T CA/09 (B).

## Supplementary Tables

**Supplementary Table 1.** H1 HA influenza virus strains and accession numbers.

| Reference Strain (Abbreviations*) | HA Accession † | Lineage/Clade |
| --- | --- | --- |
| A/South Carolina/1/1918 (SC/18) | EPI5571 | sH1N1** |
| A/Wilson-Smith/1933 (WS/33) | EPI230631 | sH1N1 |
| A/Fort Monmouth/1/1947 (FM/47) | EPI30336 | sH1N1 |
| A/Albany/12/1951 (AL/51) | EPI61365 | sH1N1 |
| A/Denver/1/1957 (DE/57) | EPI230557 | sH1N1 |
| A/USSR/90/1977 (USSR/77) | EPI390455 | sH1N1 |
| A/Brazil/11/1978 (BZ/78) | EPI57802 | sH1N1 |
| A/Chile/1/1983 (CH/83) | EPI387662 | sH1N1 |
| A/Texas/36/1991 (TX/91) | EPI18469 | sH1N1 |
| A/New Caledonia/20/1999 (NC/99) | EPI159392 | sH1N1 |
| A/Solomon Islands/3/2006 (SI/06) | EPI123307 | sH1N1 |
| A/California/04/2009 (CA/09) ‡ | EPI1859607 | pH1N1*** |
| A/New York/18/2009 (NY/09) | EPI177400 | pH1N1*** |
| A/Florida/62/2014 (FL/14) | EPI552445 | 6B |
| A/Michigan/45/2015 (MI/15) | EPI1381204 | 6B.1 |
| A/Brisbane/02/2018 (BR/18) | EPI1504919 | 1 |
| A/Hawaii/70/2019 (HI/19) | EPI1617983 | 5a.1 |
| A/Wisconsin/588/2019 (WI/19) | EPI1661231 | 5a.2 |
| A/Wisconsin/67/2022 (WI/22) | EPI2224788 | 5a.2a.1 |
| A/Netherlands/11711/2022 (NE/22) | EPI2137476 | 5a.2a.1 |
| A/swine/USA/1976/1931 (sw/USA/31) | EPI242636 | 1A.1 |
| A/New Jersey/1976 (NJ/76v) | EPI241033 | 1A.1 |
| A/swine/Hong Kong/1937/1994 (sw/HK/94) | EPI311069 | 1A.1 |
| A/swine/Ohio/891/2001 (sw/OH/01) | EPI9337 | 1A.3.3.3 |
| A/Ohio/09/2015 (OH/15v) | EPI587643 | 1A.3.3.3 |
| A/swine/England/063782/2013 (sw/ENG/13) | EPI640845 | 1B.1.1 |
| A/swine/Minnesota/A01567011/2014 (sw/MN/14) | EPI573344 | 1B.2.2.1 |
| A/northern shoveler/California/K168/2005 (ns/CA/05) | EPI230746 | n.a.‡‡ |
| A/duck/Fujian/FJ1239/2014 (dk/FJ/14) | EPI703455 | n.a.‡‡ |
| A/mallard duck/Chile/C4079/2015 (md/CHI/15) | EPI965548 | n.a.‡‡ |
| A/swine/Jiangsu/J004/2018 (sw/JS/18) | EPI1751408 | 1C.2.3 |

\* rHA generated from all H1 isolates have Y98F to reduce non-specific sialic acid binding
\*\* sH1N1: seasonal H1N1, derived from 1918 H1N1 virus
\*\*\* pH1N1: 2009 pandemic H1N1
† EPI: Epidemic Preparedness Initiative
‡ Single mutation at HA2 E47K to improve stability
‡‡ n.a.: not applicable

**Supplementary Table 2.**
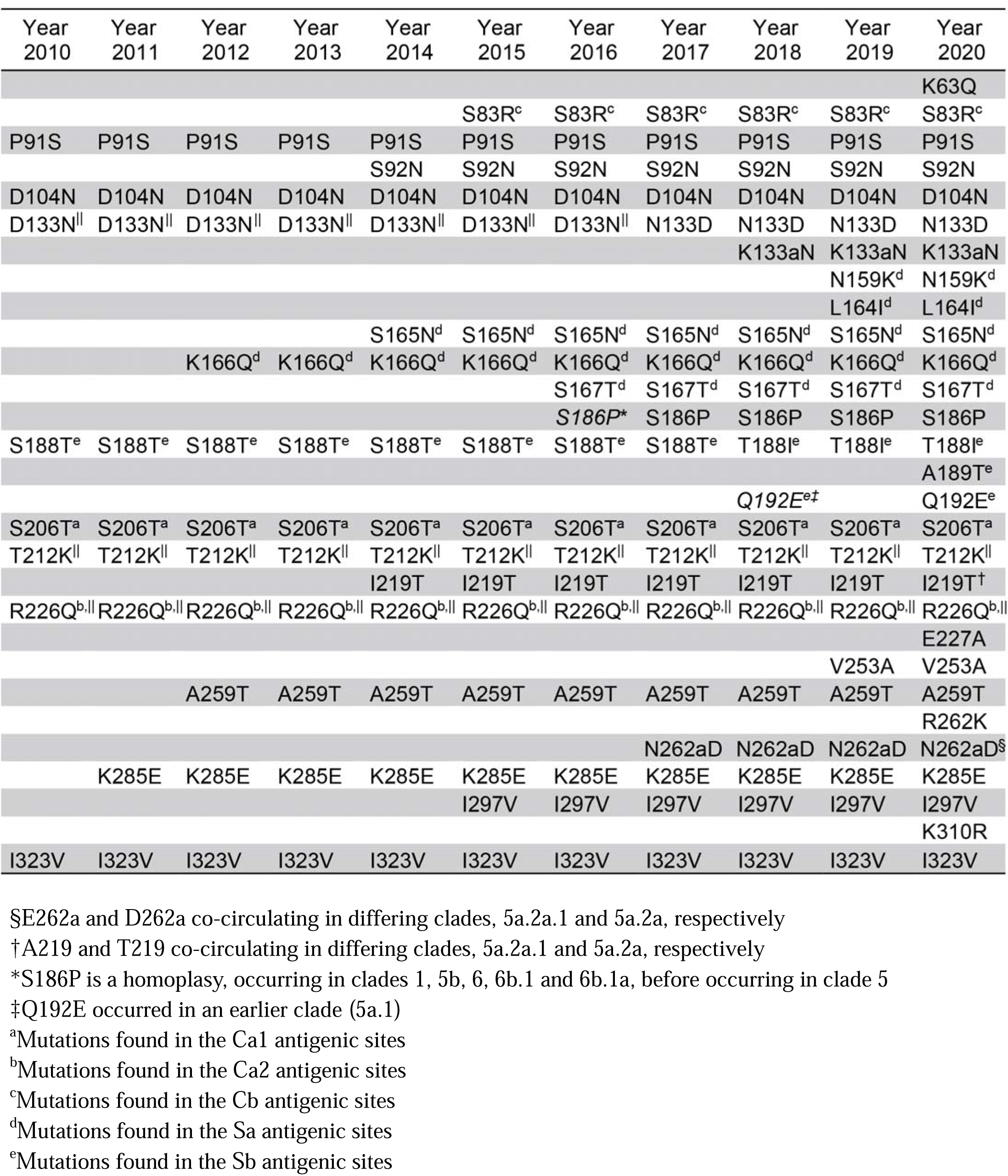
Clade-defining mutations in the H1 HA head domain from the 2010-2020. List of amino acid mutations based on Nextstrain analysis. Mutations were identified by comparing the post-2009 H1 HA isolates listed in Table S1 with the vaccine-selected reference strain A/California/07/2009.

**Supplementary Table 3:** MAbs identification, gene usage, and origin.

| Subject ID | MAb ID | Clone | Year | VH | VK/VL | Reference |
| --- | --- | --- | --- | --- | --- | --- |
| 047-09M | 2F03 | 1 | 1987 | 3-23 | K1-33 | 12, 54 |
| 047-09 | 1A02 | 1 | 1987 | 3-23 | K1-33 | 12, 54 |
| 047-09 | 4G02 | 1 | 1987 | 3-23 | K1-33 | 12, 54 |
| 047-09 | 4B06 | - | 1987 | 3-23 | K1-33 | 12, 54 |
| 240-14 | IgG 2A04 | - | 1987 | 3-23 | K1-33 | 12, 53, 54 |
| 045-09 | 2B05 | - | 1985 | 3-23 | K1-33 | 12, 54 |
| 045-09 | 2E02 | - | 1985 | 3-23 | K1-5 | 12, 54 |
| 008-10 | 5G05 | - | 1984 | 3-23 | K1-39 | 12, 54 |
| 039-10 | 5E03 | - | 1985 | 3-23 | K3-11 | 12, 54 |
| - | Ab6649 | - | 1975 | 4-39 | L1-40 | 14 |
| SFV015 | 1B03 | 2 | 1961 | 1-2 | K1-33 | 12, 54 |
| SFV015 | 1D05 | 2 | 1961 | 1-2 | K1-33 | 12, 54 |
| SFV015 | 2B04 | 2 | 1961 | 1-2 | K1-33 | 12, 54 |
| SFV015 | 2C06 | 2 | 1961 | 1-2 | K1-33 | 12, 54 |
| SFV015 | 2D02 | 2 | 1961 | 1-2 | K1-33 | 12, 54 |
| SFV015 | 2E01 | 2 | 1961 | 1-2 | K1-33 | 12, 54 |
| SFV015 | 2F02 | 2 | 1961 | 1-2 | K1-33 | 12, 54 |
| SFV015 | 3D05 | 2 | 1961 | 1-2 | K1-33 | 12, 54 |
| SFV015 | 3F01 | 2 | 1961 | 1-2 | K1-33 | 12, 54 |
| SFV015 | 3F04 | 2 | 1961 | 1-2 | K1-33 | 12, 54 |
| SFV015 | 2A06 | 3 | 1961 | 1-2 | K1-33 | 12, 54 |
| SFV015 | 2F04 | 3 | 1961 | 1-2 | K1-33 | 12, 54 |
| SFV015 | 2G04 | 3 | 1961 | 1-2 | K1-33 | 12, 54 |
| SFV019 | 1B04 | 4 | 1961 | 3-74 | K1-33 | 12, 54 |
| SFV019 | 1C05 | 4 | 1961 | 3-74 | K1-33 | 12, 54 |
| SFV019 | 2A06 | 4 | 1961 | 3-74 | K1-33 | 12, 54 |
| SFV019 | 4D05 | 4 | 1961 | 3-74 | K1-33 | 12, 54 |
| SFV019 | 4G05 | 4 | 1961 | 3-74 | K1-33 | 12, 54 |

**Supplementary Table 4:** Cryo-EM data collections, refinement and validation statistics.

| **Map** | **2F03 Fab**<br>**+ HA H1**<br>**(CA/09+E47K)** | **1A02 Fab**<br>**+ HA H1**<br>**(CA/09+E47K)** |
| --- | --- | --- |
| EMDB | 76461 | 76462 |
| PDB | 12IU | 12IV |
| **Data Collection** |  |  |
| Microscope | TFS Glacios | TFS Glacios |
| Voltage (kV) | 200 | 200 |
| Detector | Falcon 4 | Falcon 4 |
| Recording mode | Counting | Counting |
| Nominal magnification | 190,000x | 190,000x |
| EM data processing |  |  |
| Number of movie micrographs | 5616 | 5082 |
| Number of particle images in map | 135,824 | 87,530 |
| Symmetry | C1 | C3 |
| Map resolution (FSC 0.143; Å) | 3.06 | 3.7 |
| **Structure building and validation** |  |  |
| Number of atoms in deposited model | 17,160 | 13,332 |
| Residues | 2154 |  |
| Glycan | 18 | 14 |
| MolProbity score | 1.45 | 1.67 |
| Clashscore | 7.05 | 9.06 |
| Map correlation coefficient, CC (Mask) | 0.79 | 0.77 |
| d FSC model (0.5; Å) | 3.7 | 4.1 |
| RMSD Bonds |  |  |
| Bond lengths [Å] | 0.005 | 0.005 |
| Bond angles [°] | 0.966 | 0.992 |
| Ramachandran plot |  |  |
| Favored (%) | 97.75 | 97.03 |
| Allowed (%) | 2.11 | 2.97 |
| Outliers (%) | 0.14 | 0.00 |
| Side chain rotamer outliers (%) | 0 | 0 |
| cβ outlier (%) | 0 | 0 |

